# Fly Viral Atlas: A single-nucleus transcriptomic atlas of RNA viruses and transposable elements (TEs) in *Drosophila melanogaster*

**DOI:** 10.64898/2026.06.28.735102

**Authors:** Nilanjan Roy, Robert L. Unckless

## Abstract

Drosophila RNA viruses often persist in wild and lab populations, yet their tissue and cellular tropism is poorly understood. In the Fly Cell Atlas (a comprehensive Drosophila single-nucleus transcriptome) data, we detected four RNA virus infections: Nora virus, Drosophila A virus, Drosophila C virus, and Newfield virus. Nora and Drosophila A virus were the most abundant and widespread across tissues and cell types, while Drosophila C virus and Newfield virus RNA transcript were only found in oenocyte and fat body tissues. We found transcriptional changes associated with viral infection in canonical viral immunity genes (e.g. *Vago, vir-1*). Additionally, we observed that during persistent viral infections, transposable element (TE) transcripts were upregulated in somatic cells. TEs are traditionally associated with the germline, but recent studies and our data suggest they are also expressed in somatic cells. Using the Fly Cell Atlas data, we found that distinct somatic cell types express specific TE subtypes, indicating regulated and cell-type specific TE activity often overlooked in transcriptomic studies. We present Fly Viral Atlas (https://flyviralatlas.shinyapps.io/home/), a single-nucleus level atlas of RNA viruses and TE expressions in Drosophila, providing new insights into viral tropism and TE dynamics across cell types and tissues.

## 1 Introduction

Viral tropism, the selective infection of particular tissues and cell types by a virus, is a central determinant of disease outcome, transmission, and host response [1, 2]. Even within a single host, cell types vary drastically in their permissiveness to infection and in the antiviral programs they activate [3, 4, 5]. For example, Hepatitis B virus replication has been detected across diverse cell types during chronic infection, Dengue virus infection varies with host cell type and viral strain, and antiviral responses in the central nervous system differ across resident cell populations [6, 7, 8]. Defining viral tropism at cell-type resolution is therefore critical for understanding how viruses persist, disseminate, and alter host physiology.

*Drosophila melanogaster* are host to a variety of natural RNA viruses including Nora virus (NV), Drosophila A virus (DAV), Drosophila C virus (DCV), and Newfield virus (NFV). They can cause persistent infections in wild and laboratory populations[9, 10]. Nora virus is a picorna-like virus with a single-stranded positive-sense RNA genome (∼ 12 kb) and 4 open reading frames (ORFs) [11]. Interestingly, the VP1 protein encoded by Nora virus ORF1 is an RNAi suppressor[12]. Nora virus infects *Drosophila melanogaster* and related drosophilids, establishing persistent infections with minimal pathology under laboratory conditions, but notable effects on host gene expression [13, 14]. It transmits via feces (horizontal transmission, but is effectively vertically transmitted in the lab) with high pervasiveness in both laboratory and wild populations [15]. Viral titers varies (between 10^4^ and 10^10^ viral genome copies per fly in various stocks), and can persist in the lab stocks for generations [11]. The virus can evade immune pathways such as the RNAi and Toll pathway [16, 17, 18]. In terms of fitness effects, Nora virus can reduce survival, decrease offspring number, impair locomotor abilities, and increase microbiome diversity and richness in the gut [19, 20, 21, 22]. The magnitude of these effects varies by host genotype, viral load, and the abiotic factors [11].

Drosophila A virus is another persistent picorna-like virus with a ∼ 4 kb ssRNA(+) genome and 2 ORFs [23]. Drosophila A virus activates the cGAS/Sting pathway in the Drosophila gut [24], and can cause persistent viral infection. It affects lifespan, fertility, and induces immune response signaling (IMD-Relish, Sting-Relish, EGFR, JAK-STAT, JNK) [24, 25].

Drosophila C virus is a dicistrovirus with a ∼ 9 kb ssRNA(+) genome and 2 ORFs [26]. Gupta et al., 2017 intensively studied the physiological effects of Drosophila C virus in Drosophila [27]. Flies systemically infected with sublethal doses of Drosophila C virus showed increased reproductive output. In oral infections, the effect of reproductive output depends on the fly’s genetic background. Drosophila C virus infection is associated with a reduction in fecal excretion, suggesting the virus causes intestinal obstruction. Males showed a significantly greater reduction in defecation compared to females (dependent on genetic background). Survival was unaffected at very low doses (10^2^ and 10^3^ IU/mL, infectious Units per millilitre). Mortality only became appreciable at the 10^5^ IU/mL dose. Despite the lack of high mortality, Drosophila C virus can establish and replicate within the flies and higher viral titres are found in males. In terms of immunity, Drosophila C virus is constrained by the RNA interference (RNAi) pathway (by *Dcr-2 and Ago2*), along with contributions from the JAK/STAT pathway, *Toll-9* gene, and the IMD pathway [25, 28, 29, 30, 31].

Newfield virus is an unclassified member of the Permutotetraviridae with a ∼ 4.7 kb ssRNA(+) genome and 2 ORFs [32]. This virus remains understudied, and little is known about its effects on fly physiology, and the host’s transcriptomic and immune response during infection.

Studies suggest that the Nora virus infects primarily the gut (replicates in intestinal stem cells) when transmitted orally [33, 34]. The predominant infection site of Drosophila A virus, Drosophila C virus, and Newfield virus is also the gut [23, 32, 35]. However, in systemic infection, Nora virus can be detected in other tissues such as reproductive tract [36]. Subsequent work showed that Nora components circulate in the hemolymph, likely via hemocytes, and demonstrated that low-level viral RNA can be detected broadly across tissues [12, 22]. This allows spread of the virus beyond the primary site of replication (gut) [12]. This phenomenon may be associated with virus dissemination rather than true tissue infection. Nevertheless, virus dissemination in other tissues from the initial infection site is associated with upregulation of immune signaling genes and effectors including antimicrobial peptides (AMPs) [37, 38, 39, 40].

Beside influencing host gene expression, viruses can also perturb transposable element (TE) expression [41, 42, 43]. Most of the time, we associate TEs with germline expression, but recent studies have shown that they can be expressed in the somatic cells as well (somatic cells appear to control TEs through piRNAs in the fat body and through RNAi in most somatic cells) [44, 45, 46]. However, TE expression in somatic cells is poorly understood, as is there interaction with viral infection.

Interestingly, researchers frequently detect persistent RNA virus infections (viral RNA) in publicly available RNA-seq datasets (across different tissues beyond the gut), and the cell-type tropism of these viruses remains largely unknown [47]. In this study, we focus on four natural Drosophila RNA viruses that occur in the Fly Cell Atlas data: Nora virus, Drosophila A virus, Drosophila C virus, and Newfield virus. We detected these viruses (viral RNA) across different tissue samples and in the whole body [48]. Although studies show that the gut is the primary site of infection for many of these viruses, we hypothesize that as the gut barrier becomes disrupted, they spread to other tissues through the hemolymph. By utilizing the Fly Cell Atlas dataset, we aim to study the cellular tropism of persistent virus infection and associated transcriptional changes (both in terms of gene and TE expression) across different tissues. We created the Fly Viral Atlas (https://flyviralatlas.shinyapps.io/home/), a resource for exploring the cellular landscapes of viral infection and TE activity.

## 2 Materials and Methods

### 2.1 Dataset structure

To study different persistent virus infection tropism in different tissues and cells, and transcriptional changes associated with viral infections at single-nucleus resolution, we leveraged the publicly available Fly Cell Atlas data [48]. The atlas encompasses several distinct adult tissues such as head, body, antenna, haltere, proboscis and maxillary palp, wing, leg, gut, body wall, heart, testis, ovary, malpighian tubule, fat body, oenocyte, and trachea. The primary genotype used across most tissues was w1118 (*D. melanogaster*). For tissues requiring nuclear labeling to enable fluorescence activated cell sorting (FACS) based enrichment, tissue-specific GAL4-UAS transgenic lines were used: *Cg-GAL4 > UAS-lamGFP* for fat body; *drip-GAL4 > UAS-nlsGFP* for malpighian tubule; *PromE800-GAL4 > UAS-unc84GFP* for oenocyte; *btl-GAL4, tub-Gal80ts > UAS-lamGFP* for trachea.

**Table 1:** Dissection laboratories that contributed adult tissue samples to the Fly Cell Atlas dataset.

| Laboratory | Institution | Dissected tissues |
| --- | --- | --- |
| Luo lab | Stanford University | Head, body, antenna, haltere |
| O'Brien lab | Stanford University | Gut |
| Perrimon lab | Harvard Medical School | Malpighian tubule |
| Jan lab | UCSF | Body wall |
| Bodmer lab | Sanford Burnham Prebys | Heart |
| Wolfner lab | Cornell University | Male reproductive glands |
| Scott lab | UC Berkeley | Leg, wing, proboscis, maxillary palp |
| Jasper lab | Buck Institute | Fat body, oenocyte |
| Kornberg lab | UCSF | Trachea |
| Fuller lab | Stanford University | Testis |
| Nystul lab | UCSF | Ovary |
| Brueckner lab | UCSF | Hemocytes |

All tissues were dissected in cold Schneider’s medium, flash-frozen in liquid nitrogen, stored at −80°C, and shipped to Stanford University for downstream workflow.

For most tissues, 5-day-old adult flies were used. Flies were collected on day 1, kept together for mating, and on day 5 were sexed and dissected separately as males and females. There were some exceptions such as testis tissue was collected from 0–1-day old males to avoid overrepresentation of condensed haploid spermatid nuclei. Single-nucleus suspensions were prepared by homogenizing tissues. Sequencing libraries were prepared using the 10x Genomics Chromium 3*^′^* v3.1 kit and sequenced on the Illumina NovaSeq 6000. Most tissues were targeted at one 10x Genomics run per sex, with the exception of head and body, which required 6 runs each per sex given their cellular complexity, and testis, which required 3 runs to adequately capture all germ cell stages. These multiple runs reflect the sequencing depth required to reach target nucleus numbers rather than independent biological replicates. The original study also used full length RNA-seq from single cells using Smart-seq2 but in this study we only focused on the 10x samples. Refer to the Fly Cell Atlas project for more details of the samples.

### 2.2 Identification of viruses in the Fly Cell Atlas snRNA-seq data

To assess whether persistent viral infections were present in the Fly Cell Atlas snRNA-seq samples, we first downloaded all known Drosophila virus sequences from the Obbard lab (obbard.bio.ed.ac.uk/data/Updated_Drosophila_Viruses.fas.gz) [10, 47, 49]. We then aligned the snRNA-seq reads to these viral sequences using BWA mem (version 0.7.18-r1243-dirty) [50]. Viral reads were quantified with Samtools coverage (version 1.17) [51]. We applied a hard threshold of 100 RPM (reads per million) to classify samples as virus-positive. Specifically, if a viral sequence had at least 100 RPM, that sample was considered infected with the corresponding virus. Although this threshold does not confirm a robust infection, since it detects viral RNA, it helps reduce the likelihood of contamination or transient association.

Through this virus screening pipeline, we detected Nora virus, Drosophila A virus, Drosophila C virus, and Newfield virus reads in the different tissues. To confirm that putative viral reads were not mismapped from host genes, we performed BLASTN [52] searches of viral sequences against Drosophila genes. We found no BLASTN hit of the viral sequences against Drosophila genes even with a relatively low E-value cutoff of 10*^−^*^5^. We also attempted to map viral reads to the host genome without success. This approach allowed us to identify viruses present in the Fly Cell Atlas snRNA-seq data, and allowed us to study different persistent virus infection tropism and transcriptional changes in the host.

Next, we mapped snRNA-seq reads to a combined reference genome containing *D. melanogaster* transcripts (BDGP Release 6 + ISO1 MT/dm6) and the viral sequences that passed the threshold described above. This enabled us to examine transcriptional changes associated with viral infection across different Drosophila tissues and cell types. It is important to note that these detected viruses may replicate in the cytoplasm (their nuclear presence remains unknown and understudied) [11, 23, 26, 32]. However, we detected viral reads in snRNA-seq data, likely reflecting cytoplasmic contamination during nucleus isolation, which is common in snRNA-seq preparations [53, 54].

### 2.3 Data processing and analyses

Barcode processing along with count matrix generation was performed separately for each tissue of the Fly Cell Atlas snRNA-seq samples using CellRanger (version 7.2.0) and SoloTE (version 1.09) [55, 56]. BAM files generated by CellRanger for different tissue samples were used as input for SoloTE. SoloTE generated count matrices for genes, detected viruses, and TEs. A combined transcriptome containing the Drosophila and viral genomes (BDGP Release 6 + ISO1 MT/dm6, Nora virus; NC 007919.3, Drosophila A virus; NC 012958.1, Drosophila C virus; NC 001834.1, Newfield virus; MH384307.1) was used to quantify gene and viral counts, while repeatMasker output from BDGP Release 6 (dm6) was used to quantify TEs. SoloTE assigns reads to annotated genes only if reads map within one of their exons, then excludes gene-assigned reads to avoid falsely counting TEs located within genes as expressed. SoloTE then evaluates remaining unassigned reads for overlaps with TE annotations and quantifies TE expression at both locus and family levels.

We filtered the raw gene-barcode count matrix generated by SoloTE with DropletUtils (version 1.30.0) to omit barcodes associated with background noise such as free floating ambient mRNA from lysed or dead cells [57, 58]. Doublets were removed using scDblFinder (version 1.24.10) [59]. Then the count matrix was processed and analyzed with Seurat (version 5.5.0) [60] for each tissue separately. For creating the seurat object, we used a minimum cell count *>*3, and minimum features *>*200. We merged each replicates to do inegrated downstream analyses for each tissue samples individually. We used Harmony (version 2.0.3) for batch correction and integration of replicates of each tissue samples [61]. Cells with 200 to 2500 genes, and *<*25% mitochondrial genes were retained. This filtering removed low quality cells and empty droplets, which typically contain very few detected genes, as well as potential doublets with unusually high gene counts. Scaling and log normalization were performed using standard Seurat functions with default parameters. The top 40 PCs were selected based on the elbow plot and were used for principal component analysis (PCA) and uniform manifold approximation and projection (UMAP). Cell type clusters were identified at a resolution of 0.5. Cell types were identified using marker genes from the Fly Cell Atlas project, and ScType was used to automate the process [62]. To measure viral and TE expression (RNA) tissue-wide, cells were pooled by sex, and to measure expression by cell type, cells were pooled by the combination of cell type and sex. Within each pool, we calculated counts per million (CPM) by summing raw counts across cells and scaling to one million. Thus, we calculated the expression (RNA) of Nora virus, Drosophila A virus, Drosophila C virus, Newfield virus, and TE subfamilies in different tissue samples and cell types. We also analyzed ambient RNA corrected counts generated with decontX (version 1.8.0) to compare these estimates with our SoloTE derived counts [63].

To identify differentially expressed genes and TEs associated with viral infections in different tissues and cell types, we performed differential expression analyses with the seurat FindMarkers function. We used Seurat MAST model for differential expression testing as it allowed to control for covariates such as cell types, replicates, sex, other virus infections (e.g., controlling Drosophila A virus infected cells while doing differential expression testing for Nora virus), and interaction effects (e.g., cell types and virus interaction, interactions between viruses). Before doing the differentially expressed genes analyses, we defined cells as virus infected if the cell had viral titer (normalized count) greater than 0. We performed differential expression analysis at two levels: tissue-wide, pooling all cells, and within each annotated cell types. At the tissue-wide level, for each virus of interest we fit the model: gene/TE expression ∼ virus*_i_*+cell type+virus*_j_* +sex+virus*_i_* : cell type+virus*_i_* : virus*_j_*, where virus*_i_* denotes the focal virus being tested and virus*_j_* denotes co-infecting virus(es) included as covariates, with infection status for each virus coded as 0 (uninfected cells) or 1 (infected cells). We defined cells as virus infected if the cell had viral titer (normalized count) greater than 0. Using this approach, we labelled each cell according to its infection status with specific viruses. The focal virus virus*_i_* was treated as the variable of interest, while cell type, co-infection status, sex, the virus–cell type interaction, and the virus–virus interaction were included as latent variables to control for cellular composition, co-infection, and sex-specific effects. To identify cell-type specific responses, we further subset the data by annotated cell type and fit for each virus of interest within each cell type. The model we used: gene/TE expression ∼ virus*_i_* +virus*_j_* +sex +virus*_i_* : virus*_j_*. Cell type was not included as a covariate in this stratified model because each test was performed within a single cell type. Cell types with fewer than 30 cells in either infection class were excluded to ensure sufficient statistical power. The expression of piRNA pathway genes in the somatic tissue is generally very low (they are mostly active in germline cells) [25, 64]. For this reason, mean counts of piRNA pathway genes were calculated from only cells that had piRNA pathway gene expression and were measured at the tissue level. However, we downsampled the total number of cells to an equal number of cells for the uninfected and virus infected cells categories before calculating mean counts of piRNA pathway genes. This was done to randomly select cells that expressed piRNA pathway genes. Without downsampling, we found that piRNA genes are mostly upregulated in virus infected cells, because the number of virus infected cells is low compared to uninfected cells. Thus, mean counts of piRNA are most of the time high in virus infected cells compared to the uninfected cells.

Pathway enrichment analysis was conducted with Flyenricher [65, 66]. R (version 4.5.3) [67] and R packages tidyverse (version 2.0.0) [68] and shiny (version 1.13.0) [69] were used for data wrangling, plotting and making web applications. Additional figure editing was performed using Microsoft PowerPoint.

## 3 Results

### 3.1 Drosophila RNA viruses and TEs exhibit distinct tissue level tropism

To understand tissue wide viral and TE tropism in Drosophila, we examined the normalized counts (CPM) across tissues and cell types. For viruses, we considered counts as proxy for viral titer (RNA), while TE counts represents transcript abundance.

RNA from Nora virus and Drosophila A virus was detected in nearly all examined tissues (Figure 1a). We hypothesize that these viruses may disseminate through the hemolymph from disrupted gut (the primary infection site) and spread to other tissue [12]. In some tissues, such as the body wall, Nora virus titer was more abundant than Drosophila A virus but that was only observed in female, while the opposite pattern (Drosophila A virus more abundant than Nora virus) was observed in the fat body (both males and females). Previous studies have shown that infection route is likely fecal-oral [22, 23, 34], so direct comparisons of viral titer between viruses are difficult and may not be reliable. In contrast, RNA from Drosophila C virus and Newfield virus was detected at appreciable levels only in the fat body and oenocyte. Notably, tissues were collected and processed by different laboratories. It is possible that Drosophila C and Newfield virus were simply absent from the fly populations in the labs that contributed other tissues. Additionally, we noticed female flies exhibited higher viral RNA levels compared to males for Nora and Drosophila C virus but not for Drosophila A and Newfield virus (although no statistical significance was found in any comparisons, wilcox test p value *>* 0.3) (Supplementary figure S1a). A previous study has shown that, after mating immune system is compromised in female flies [70].

**Figure 1:**
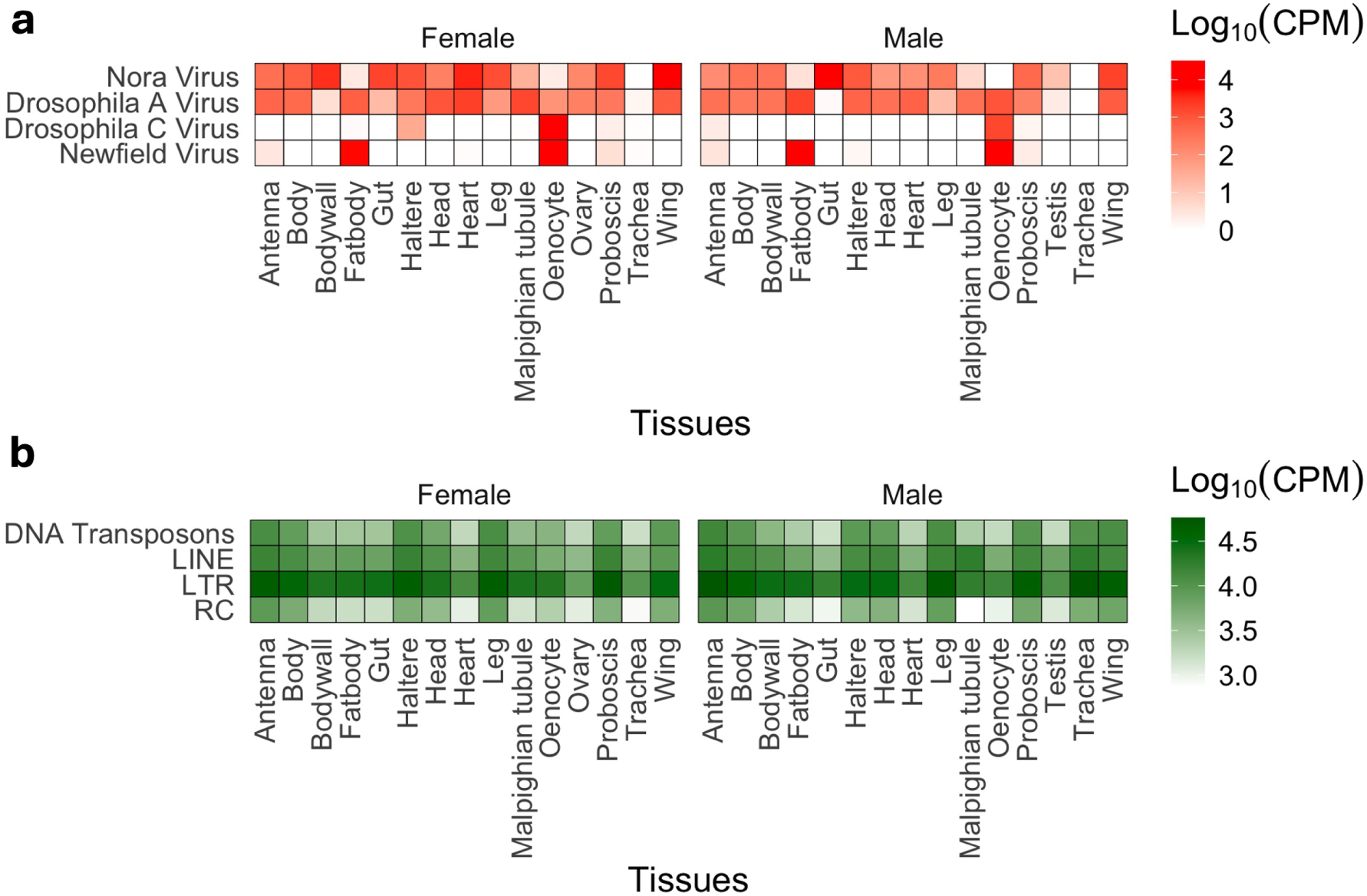
Tissue level tropism of RNA viruses and TEs in Drosophila. (a) Viral RNA read counts across different tissues of female and male flies. (b) TE transcripts across different tissues of female and male flies. 4 classes of TE transcripts were measured; DNA transposon, RC (Helitrons), LTR (retrotransposon), and LINE (retro-transposons). Raw reads were normalized to counts per million (CPM). CPM values were then log_10_ transformed to visualize the range.

Regarding TEs, we observed exceptionally high abundance of TE transcripts across somatic tissues (Figure 1b). Among four main TE classes, long terminal repeat (LTR) elements were the most abundant across tissues, likely because they are the most widely represented class of TEs in the Drosophila genomes (Figure 1b) [71]. Interestingly, several somatic tissues (antenna, haltere, and head) showed higher TE expression than germline tissues (ovary and testis), potentially because the piRNA pathway is more able to suppress expression in germ cells [64]. We also observed higher TE expression in male tracheal tissue (respiratory organ) than in female trachea.

Overall, we observed detectable levels of viral RNA across a wide range of tissues in persistent Nora and Drosophila A virus infection, but much more tissue specific infection of Drosophila C and Newfield virus. Somatic tissues also show detectable TE transcript levels, supporting the idea that somatic TE expression is biologically important.

### 3.2 Drosophila RNA viruses exhibit distinct cell tropism across tissues

We wanted to understand cell level tropism of the persistent RNA viruses in Drosophila. For that, we clustered cells from tissues based on similarity of gene expression using the Fly Cell Atlas snRNA-seq data, and annotated the clustered cells with maker genes (a subset of genes with distinct expression in specific cell types, see methods). After identifying the cell types in each tissue, we looked at the viral RNA levels (proxy for viral titer) to observe the cell tropism.

Since the whole body data encompasses cell types from all tissues (Supplementary figure S2), we first examined virus tropism in whole body context. Only Nora and Drosophila A virus were present in the whole body data (Figure 1a), and we found that they are most prevalent in cells associated with indirect flight muscles (Figure 2a). Overall, both viruses show similar cell tropism, perhaps because they belong to the picorna virus family [11, 23]. To see if this holds true for other dissected tissue, we looked at the haltere (Supplementary figure S3) which is a muscle-related tissue. We found similar cell tropism for both viruses where muscle cells and adult fat body cells were infected (Figure 2b). In the Fly Cell Atlas data, we could detect Drosophila C virus and Newfield virus only in the fat body and oenocyte (Figure 1a). In the fat body (Supplementary figure S4), Newfield virus RNA was detected across all annotated cell types. In contrast, Drosophila A virus RNA was mainly concentrated in male germline-associated and unannotated (marker genes: *Desat1, Yp2, Yp3, RpLP2, RpS29, RpL19, RpLP0, Gapdh2, CG5791, CG16836, CG16713*) cell types (Figure 2c). In the oenocyte (Supplementary figure S5), RNA reads from Drosophila A, Drosophila C and Newfield viruses were detected. Drosophila C and Drosophila A virus RNA was mostly abundant in the unannotated cell type (marker genes: *Sfp87B, Sfp60F, Desat1, Mlc2, Cyp4g1, CG13315, EbpII*), while Newfield virus RNA was broadly distributed across all oenocyte cell types (Figure 2d). We also compared the results with decontX processed counts which corrects for ambient RNA contamination, and we did not find any significant changes in tropism (Supplementary figure S6, S7, S8, S9).

**Figure 2:**
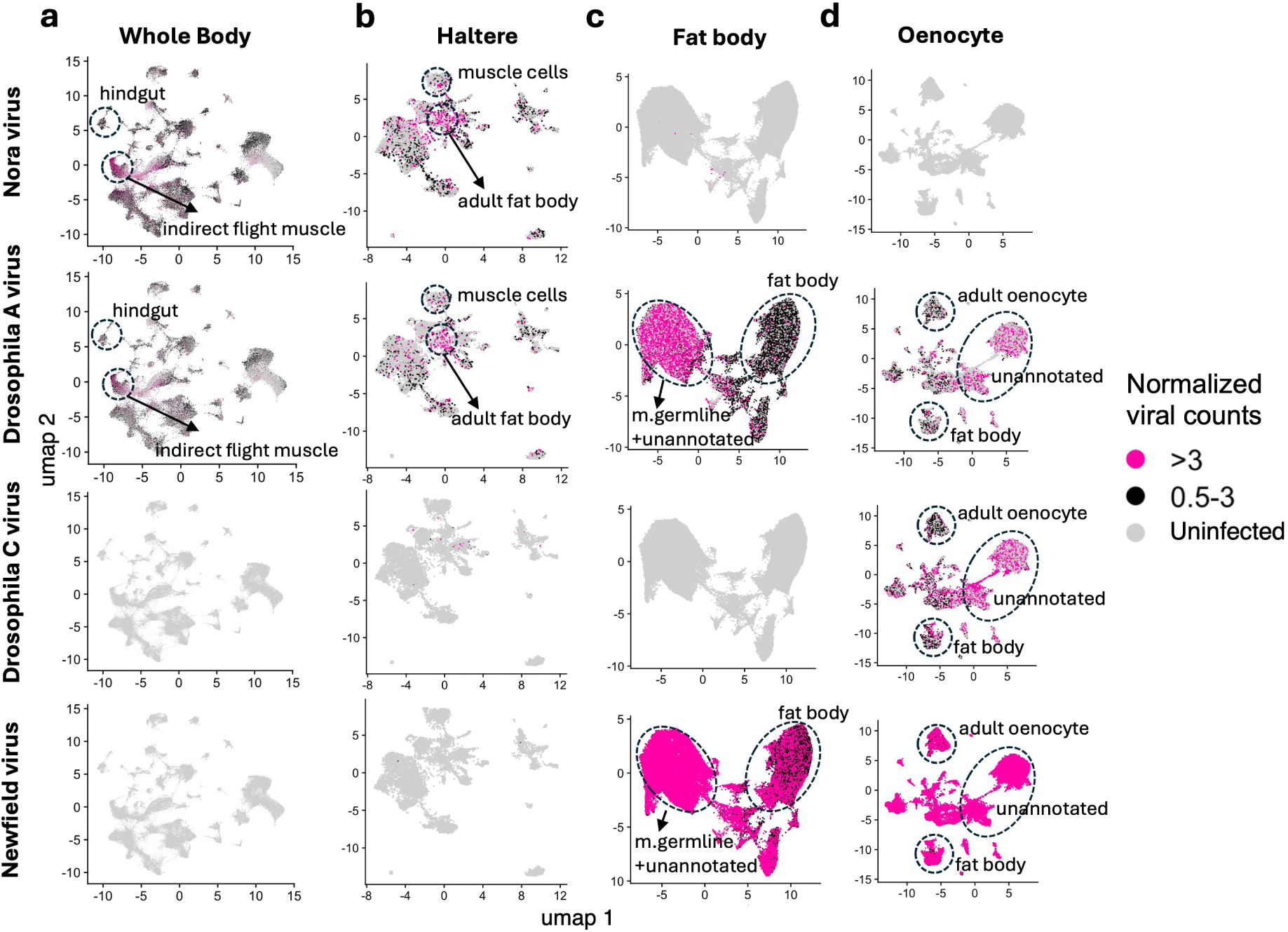
Cellular tropism of RNA viruses in Drosophila. UMAP plots were generated to visualize viral RNA reads from the whole body (a), haltere (b), fat body (c), and oenocyte (d). Counts are log normalized (which is the standard normalization method in UMAP based visualization for the single nucleus data). Seurat LogNormalize method was used where feature counts for each cell are divided by the total counts of that cell, multiplied by a scale factor of 10,000, and then taking natural log–transformed value using log1p. *>*3 represents cells where viral RNA counts are greater than 3. 0.5-3 represents cells where viral read counts are between 0.5 to 3. Uninfected represents cells that don’t have any viral RNA. Log normalized counts can be converted to CPM using the formula *CPM* = (*e*^LogNormalized^ ^count^ − 1) × 100. Thus, values *>* 3 correspond to viral RNA reads greater than ∼2000 CPM, while values between 0.5 and 3 correspond to viral RNA reads between ∼65–2000 CPM.

Overall, persistent RNA viruses may show distinct cell-type tropism rather than uniform distribution across infected tissues. Interestingly, Nora virus and Drosophila A virus showed enrichment in muscle-associated cell types.

### 3.3 Transposable elements exhibit cell type specific expression patterns in both germline and somatic cells

To investigate the cellular tropism of TEs across tissues, we analyzed TE transcript abundance across distinct cell types from multiple tissues using transcript counts generated with SoloTE. Similar to the viruses, first we investigated the whole body sample (Supplementary figure S2, figure 3a). In the whole body, *G2* elements (subclass:Jockey, class:LINE) were predominantly expressed in spermatocytes (germline cells), while *mdg4* formerly known as *gypsy* (class:LTR) was primarily expressed in epithelial (somatic) cells (Figure 3a). Similar patterns were observed in other tissues, including the testis (germline tissue) and leg (somatic tissue) (Supplementary figure S10, S11, and Figure 3b, c). However, we also found that TEs such as *DNAREP1* (Helitron:RC), *ROO* (Pao:LTR), *TABOR* (mdg4;Gypsy:LTR), *FW* (Jockey: LINE) had broad expression level in both somatic and germline tissue (Supplementary figure S12, S13, S14).

**Figure 3:**
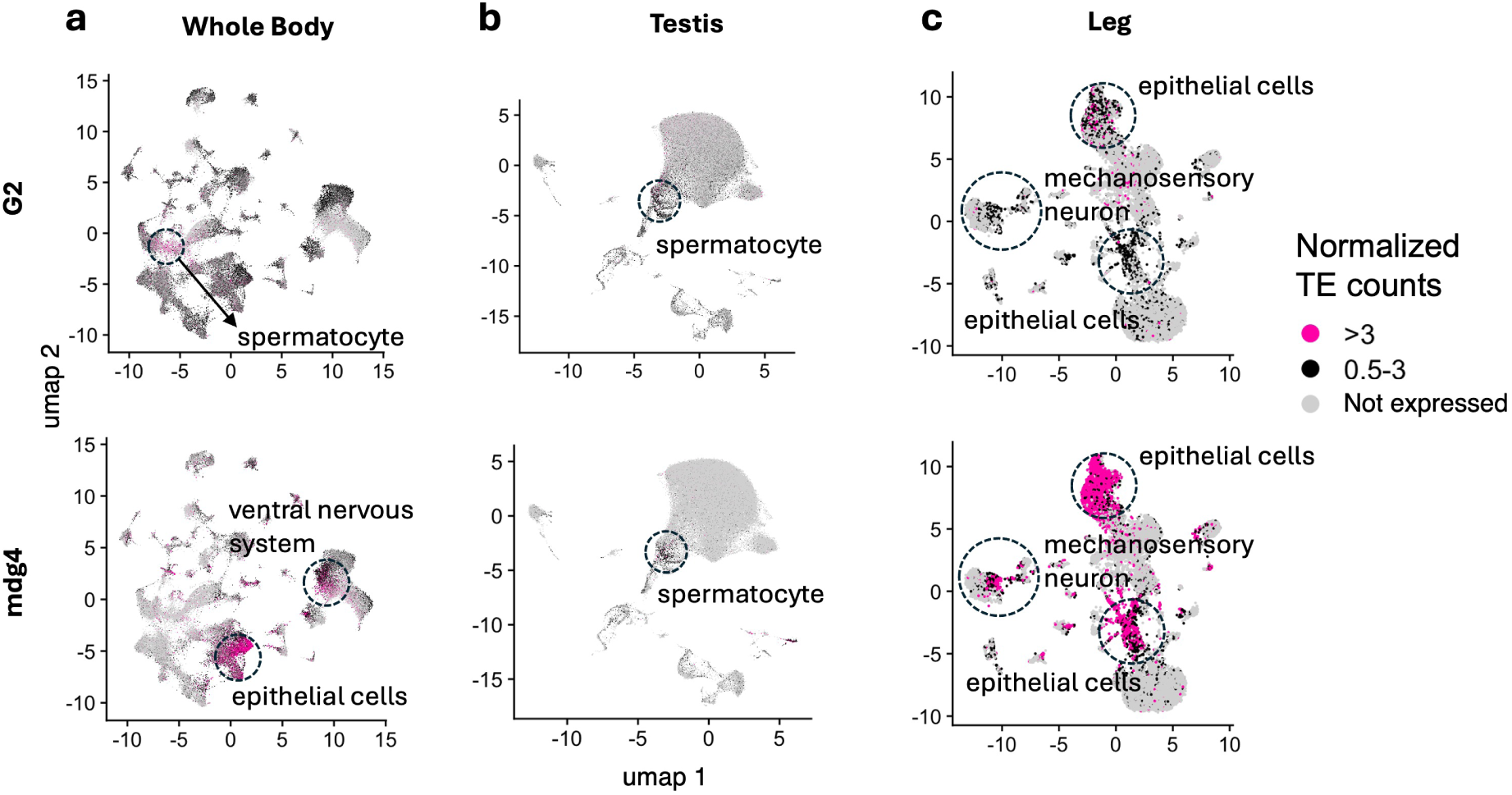
Cellular tropism of TEs in Drosophila. UMAP plots were generated to visualize TE transcripts from the whole body (a), testis (b), and leg (c). Counts are log normalized (which is the standard normalization method in UMAP based visualization for the single nucleus data), and represents TE expression. Seurat LogNormalize method was used where feature counts for each cell are divided by the total counts of that cell, multiplied by a scale factor of 10,000, and then taking natural log–transformed value using log1p. *>*3 represents cells where TE expression counts are greater than 3. 0.5-3 represents cells where TE expression counts are between 0.5 to 3. Not expressed represents cells that don’t have any TE expression. Log normalized counts can be converted to CPM using the formula *CPM* = (*e*^LogNormalized^ ^count^ − 1) × 100. Thus, values > 3 correspond to TE expression greater than ∼2000 CPM, while values between 0.5 and 3 correspond to TE expression between ∼65–2000 CPM.

Together, these results suggest that TEs exhibits strong cell type specificity across Drosophila tissues in both germline and somatic cells.

### 3.4 Persistent RNA virus infection alter host transcription

We aimed to investigate the host response to persistent RNA virus infection by utilizing the four RNA viruses identified in the Fly Cell Atlas dataset. We performed differential gene expression analysis by comparing virus infected cells with the virus uninfected cells (see methods for full model). Although, we did the analyses for each dissected tissues (accessible through the shiny web application), here we are only focusing on the results of (differential gene expression in Nora and Drosophila A virus) the whole body samples. We found canonical antiviral genes are upregulated (*AGO2, Vago, Vir-1, ref(2)P*) in association with the Nora virus infection (Figure 4a). Immune response pathways such IMD (*AttB*) and Toll (*Mtk*) are also upregulated (Figure 4a). A handful of genes such as *lncRNA:roX1* (involved in dosage compensation), *CG15120* (ubiquitin and heat shock protein binding activity), *Met75Ca/Cb* (insemination) are downregulated in association with the Nora virus infection (Figure 4a). In terms of gene ontology (GO) analysis, cytoplasmic translation, metabolic process came up as top enriched GO terms (Supplementary figure S15) associated with Nora virus infection. Drosophila A virus infection-associated differentially expressed genes largly overlap with the ones we found with Nora virus (Figure 4b). Although there were exceptions, for example, Cecropins (*CecC, CecA2*) which are involved in the systemic immune response and in various epithelia were upregulated in association with Drosophila A but not Nora virus. The majority of the GO terms associated with Drosophila A virus infection were similar to Nora virus (Supplementary figure S16). We further analyzed the gene expression patterns (showing here a handful of immune response related genes) in cell types (Figure 4c). We found that, some genes show distinct up or downregulation patterns based on cell types, in other words they show a significant tissue by infection interaction. For example, *PGRP-LC* (receptor protein in IMD pathway) shows higher infection-associated expression in most of the cell types except for the adult oenocyte. Results from additional tissues are accessible via the Shiny web application (Section 3.6). One caveat of this differential gene expression analysis is the issue of pseudo replication when comparing virus-infected cells with virus-uninfected cells. However, our earlier study showed that, even with this limitation, biological signals are detectable in terms of differentially expressed genes [25]. We also compared differential gene expression results across the same cell types derived from different tissues to assess the degree of concordance between tissues (Supplementary figure S17). We observed significant concordance in differential expression associated with infection among epithelial cell types across different tissues, indicating a shared transcriptional response, and this was not evident for dissimilar cell types, where differential expression signatures showed substantially lower similarity across tissues (as expected). However, this pattern was detected only in epithelial cells (which one of the most common cell types in tissues [72]), we suspect that the Fly Cell Atlas dataset lacks sufficient statistical power to robustly identify differentially expressed genes at the resolution of individual cell types.

**Figure 4:**
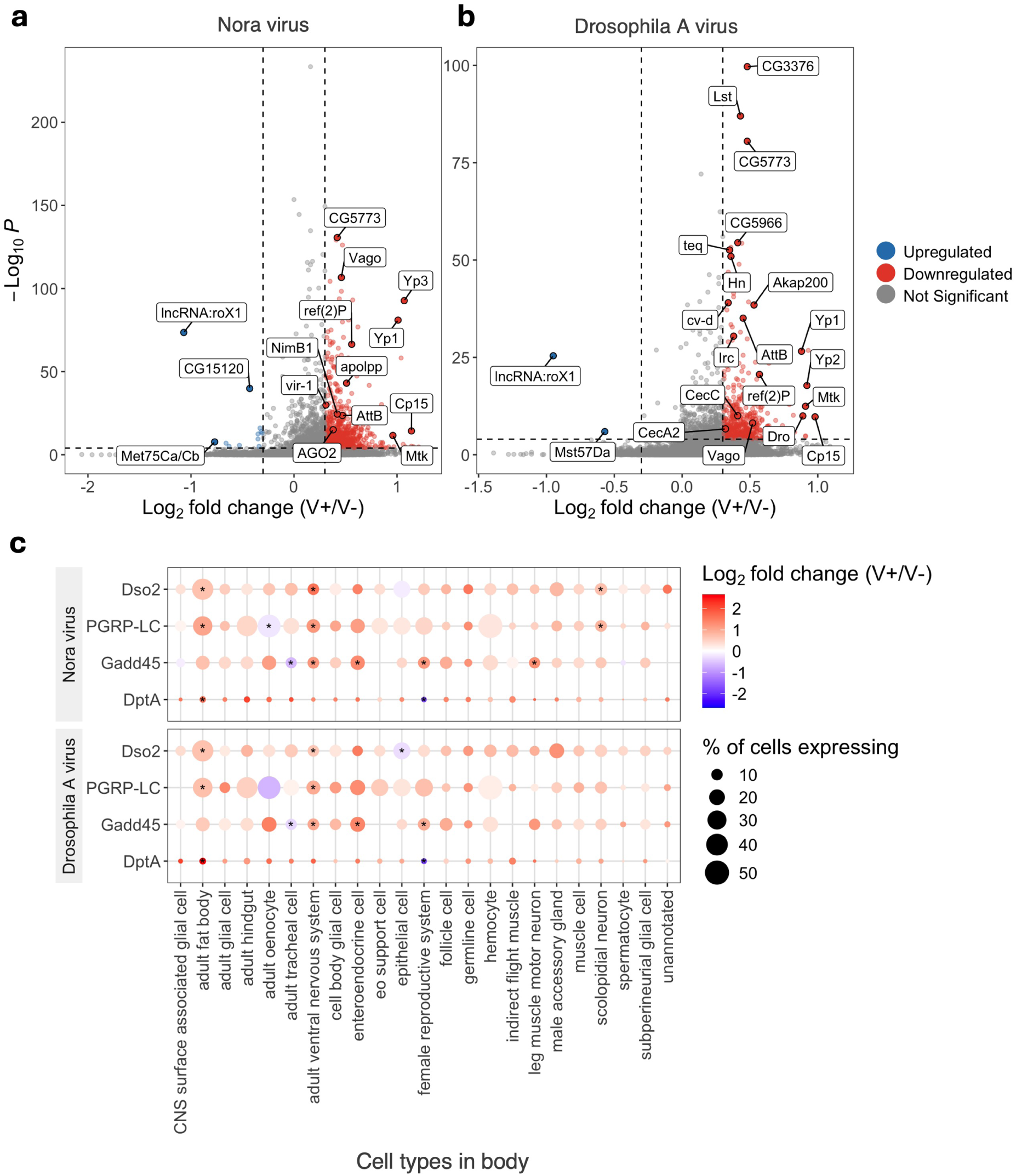
Differential gene expression analysis during viral infection in the Drosophila whole body sample. (a) Volcano plot of differentially expressed genes during Nora infection. (0.05/total number of genes) was used as *pvalue* threshold. Absolute value of 0.3 was used as threshold for Log_2_ fold change. (b) Volcano plot of differentially expressed genes during Drosophila A virus infection. The same thresholds were applied as in panel (a). (c) Cell type specific expression changes of immune related genes in viral infection in the whole body. Virus infected cells (V+) was compared against virus uninfected cells (V-). We defined cells as virus infected if the cell had viral titer (normalized count) greater than 0. Using this approach, we classified each cell based on its infection status with Nora virus or Drosophila A virus. * means statistically significant (Seurat MAST model p-adjusted value *<* 0.05).

Overall, our findings lead us to hypothesize that persistent viral infection leads to sustained elevation of immune response gene expression throughout the adult lifespan of the fly.

### 3.5 Persistent RNA virus infection upregulate transposable elements

We wanted to understand how TE expression changes in association with persistent viral infections specially in somatic cells. From the whole body data, we saw that irrespective of the TE subfamilies, TEs are upregulated in association with persistent virus infections (Figure 5a). We also looked at the piRNA (canonically germline and fat body regulation) and RNAi (canonically somatic cell regulation) pathways that regulates the TEs [44, 45, 46]. We found that genes involved in these pathways are overwhelmingly downregulated in association with persistent viral infections, which is consistent with the upregulation of TEs (Figure 5b). However, very few counts of piRNA pathway genes can be detected in somatic cells, and the snRNA-seq data we are analyzing is not made to measure such low counts. Furthermore, lowly expressed genes leads to zero inflation in snRNA-seq data [25, 73]. Thus, there is considerable amount of noise when measuring piRNA pathway genes. We found similar results (TE upregulation) when we looked at significantly (Seurat MAST model p-adjusted value *<* 0.05) differentially expressed TEs in viral infections at other dissected tissues except in proboscis where we found traffic jam (*tj*), which is a master regulator of piRNAs in somatic cells [74] is upregulated specifically in Drosophila A virus infected cells (Supplementary figure S18, S19).

**Figure 5:**
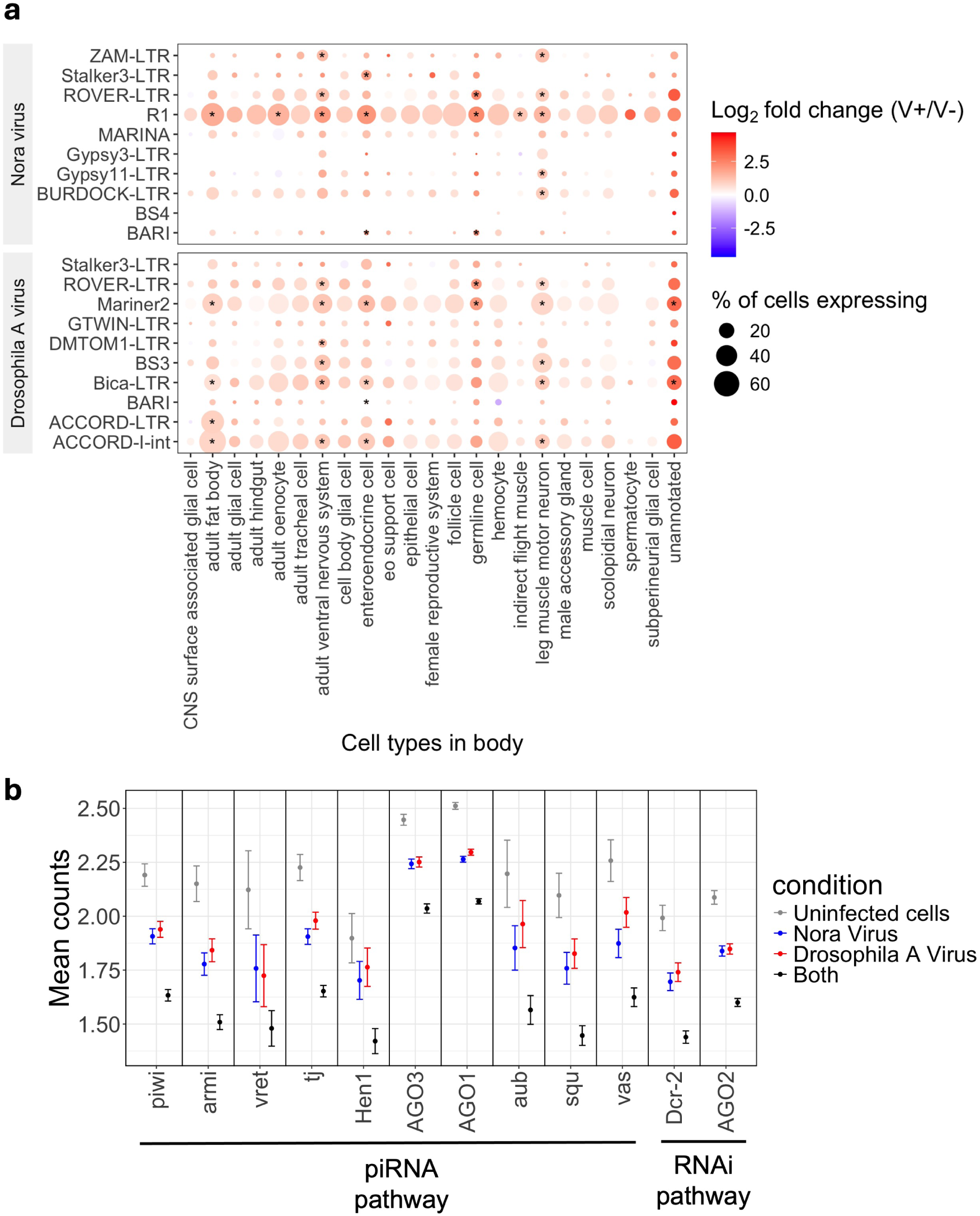
TE dynamics in persistent RNA virus infection in the Drosophila whole body. (a) Expression changes of top 10 TE subfamilies in Nora and Drosophila A virus infection. (b) Expression of piRNA and RNAi pathway genes in virus infection in the whole body sample. Uninfected cells are those with zero RNA reads for both Nora and Drosophila A viruses. Drosophila A virus infected cells contain Drosophila A virus RNA reads (*>*0) but no Nora virus reads (0). Nora virus infected cells contain Nora virus RNA reads (*>*0) but no Drosophila A virus reads (0). Both means cells have both Nora and Drosophila A virus RNA reads(*>*0). Counts are normalized with standard Seurat NormalizeData function. * means statistically significant (Seurat MAST model p-adjusted value *<* 0.05).

Overall, we hypthesize that the host’s control over TEs in both germline and somatic cells may get disrupted in persistent RNA virus infections, perhaps due to the viruses manipulation of RNAi pathways.

### 3.6 Fly Viral Atlas: Interactive shiny web application to explore viruses and transposable elements expression in cell types of adult Drosophila

In this study we analyzed persistent RNA virus titers, TE expression patterns, and differential gene and TE expression associated with viral infection, primarily focusing on whole-body samples. as mentioned earlier, all analyses were performed separately for each dissected tissue, enabling tissue-specific investigation of viruses and TEs in the host. To facilitate access and exploration of these analyses, we developed an interactive Shiny web application (https://flyviralatlas.shinyapps.io/home/) (Figure 6). The application provides users with the ability to: (1) examine RNA virus and TE read counts across cell types in adult Drosophila tissues. To do this, users can navigate to the cell type specific viral/TE reads section, where they can select a virus or TE of interest and visualize its read counts across different tissues. Alternatively, a separate cell types tab allows users to select a specific tissue and examine read counts across its constituent cell types; (2) explore differential gene and TE expression associated with four persistent RNA virus infections. In the differential gene/TE expression in viral infection section, users can select a gene or TE of interest and view its log_2_ fold change across different tissues under different viral infection conditions. A separate tab enables users to choose a specific tissue and investigate log_2_ fold changes across the cell types within that tissue; and (3) visualize the distribution and tropism of RNA viruses and TEs using UMAP-based representations. For each tissue, dedicated sections allow users to select a virus or TE of interest and explore its distribution and cell type tropism. Figure 6 shows only a representative snapshot of the application and its features.

**Figure 6:**
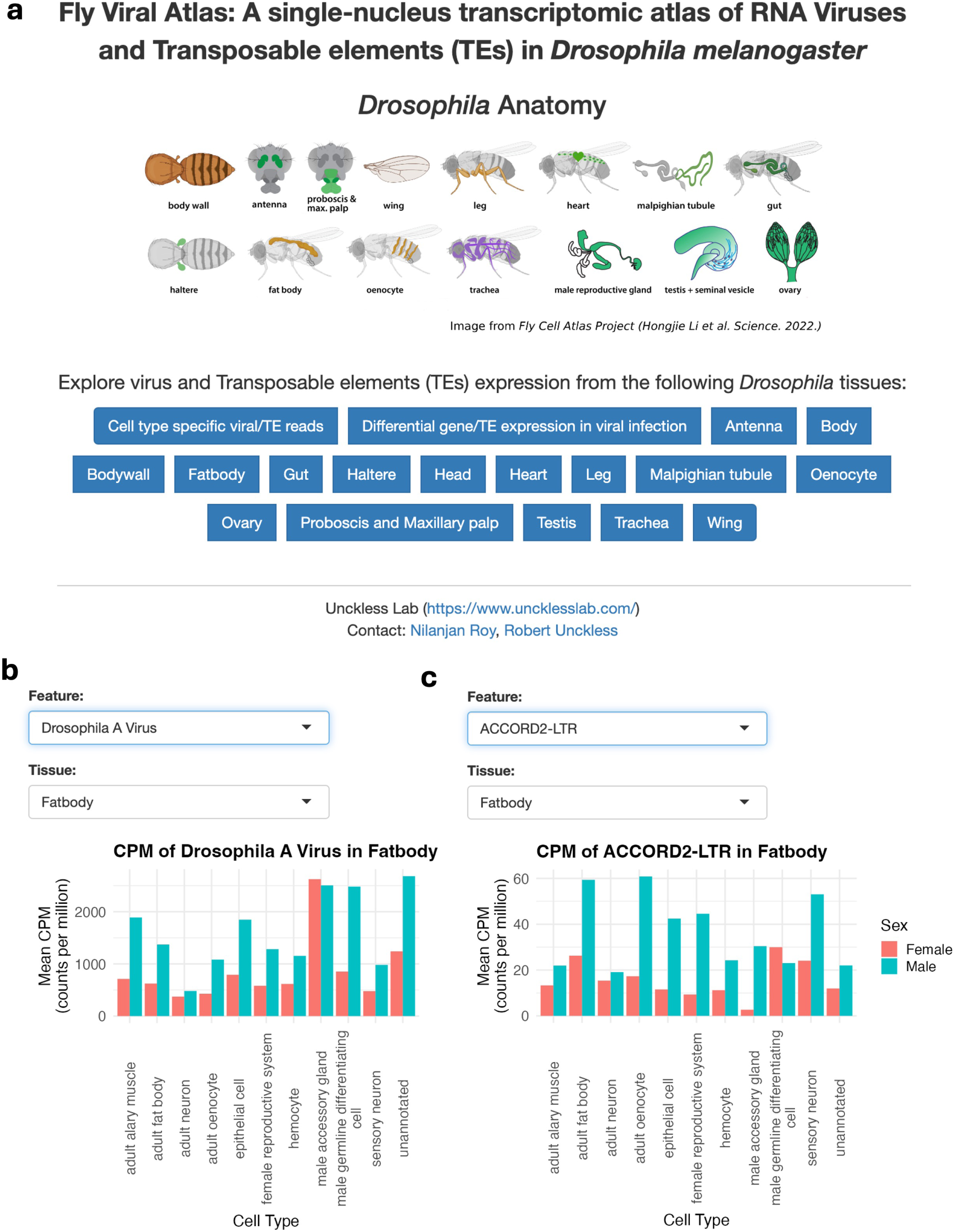
Fly Viral Atlas shiny web application for interactive exploration of RNA virus and TE expression across adult Drosophila tissues. (a) A partial snapshot of the application interface and navigation options. (b) Example visualization of RNA virus read counts across cell types and tissues, allowing users to examine viral distribution and cell-type tropism. (c) Example visualization of TE expression across cell types and tissues, enabling comparison of TE abundance and specificity across cell types and tissues. The application also provides access to differential gene and TE expression results associated with persistent RNA virus infection, along with UMAP-based visualization of the distribution and tropism of RNA viruses and TEs across cell types of tissues.

By enabling interactive exploration of the data, the web application facilitates the development of new hypotheses related to viral infection, host responses, and TE regulation.

## 4 Discussion

We studied several persistent RNA viruses and TEs in Drosophila and characterized their tissue and cell-type tropism across the fly. In corcordance with other studies, we found that Nora and Drosophila A virus are extremely common in the fly lines researchers maintain in the laboratory [34, 47]. We observed that Nora and Drosophila A virus RNA reads can be detected at single cell level across tissues although their primary site of infected is gut [22, 23]. This, along with other studies, leads us to hypothesize that when the gut barrier is disrupted, viruses or viral particles/genomes can enter the hemolymph and reach other tissues, potentially leading to infection in those tissues [12, 36]. This may be the case of other persistent viruses such Drosophila C and Newfield virus as well, but with the data that we used, we could only detect them in the fat body and oenocyte (their primary infection site is also the gut) [32, 75, 76]. A limitation is that tissues came from different laboratories, so virus absence may reflect lab-specific fly populations rather than tissue specificity.

In terms of cellular tropism, we found that Nora and Drosophila A virus exhibited the highest viral titers (viral RNA levels) in muscle-associated cells in whole body samples. This finding was unexpected, as we initially anticipated that these viruses would primarily target the gut. Nevertheless, substantial viral RNA levels were also detected in gut tissues, indicating that the gut remains an important site of infection. A previous study reported locomotor abnormalities in flies infected with Nora virus, and the higher viral titers detected in muscle-related cells in our study are consistent with this observation [21]. In terms of host response to viral infection, we found immune pathways such as IMD, Toll, antiviral genes (*Vago, vir-1*) and they are consistently associated with these type of persistent viral infections [9, 25].

We detected abundance of TE transcripts (LTR being the most abundant) in somatic tissues. High levels of somatic TE activity for the three most abundant TE families (*Alu*, *SVA*, and *L1*) across diverse tissue types (breast, head, lung) has been also noted in humans [77]. We also observed cell type specific TE expression, for example, *G2* in spermatocyte cells, mdg4 (formerly gypsy) in epithelial and neuronal cells. Some TEs like *DNAREP1, ROO* are found to have broad expression pattern in tissues. This type of pattern has been also reported in somatic tissue in other studies [44, 56]. In terms of TE regulation during viral infection, we found that TEs were predominantly upregulated in response to infection. This observation is consistent with findings from our previous study as well as reports from other studies [25, 41, 43].

Importantly, we developed the Fly Viral Atlas (https://flyviralatlas.shinyapps.io/home/) web application to provide an interactive platform for exploring RNA virus and TE analyses generated in this study. The application enables users to explore viral and TE read counts, cell tropism, and differential gene expression across adult Drosophila tissues and cell types. We hope that the application will be a useful resource for insect virologists and TE researchers community.

## Acknowledgements

We thank members of the Unckless lab for their valuable feedback.

## Author information

**Department of Molecular Biosciences, The University of Kansas, USA.**

Nilanjan Roy, Robert L. Unckless

## Contributions

Conceptualization: NR, RLU; Data curation: NR, RLU; Formal analysis: NR, RLU; Funding acquisition: RLU; Investigation: NR, RLU; Methodology: NR, RLU; Project administration: NR, RLU; Visualization: NR, RLU; Writing: NR, RLU.

## Ethics declarations

## Ethics approval and consent to participate

Not applicable

## Consent for publication

Not applicable

## Supplementary information

**Additional file, SM1:** Supplementary figures.

## Availability of data and material

The processed data and code will be made available in the peer-reviewed publication.

