## Supplementary Materials for "Fly Viral Atlas: A single-nucleus transcriptomic atlas of RNA viruses and transposable elements (TEs) in *Drosophila melanogaster*"

### 1 Additional figures

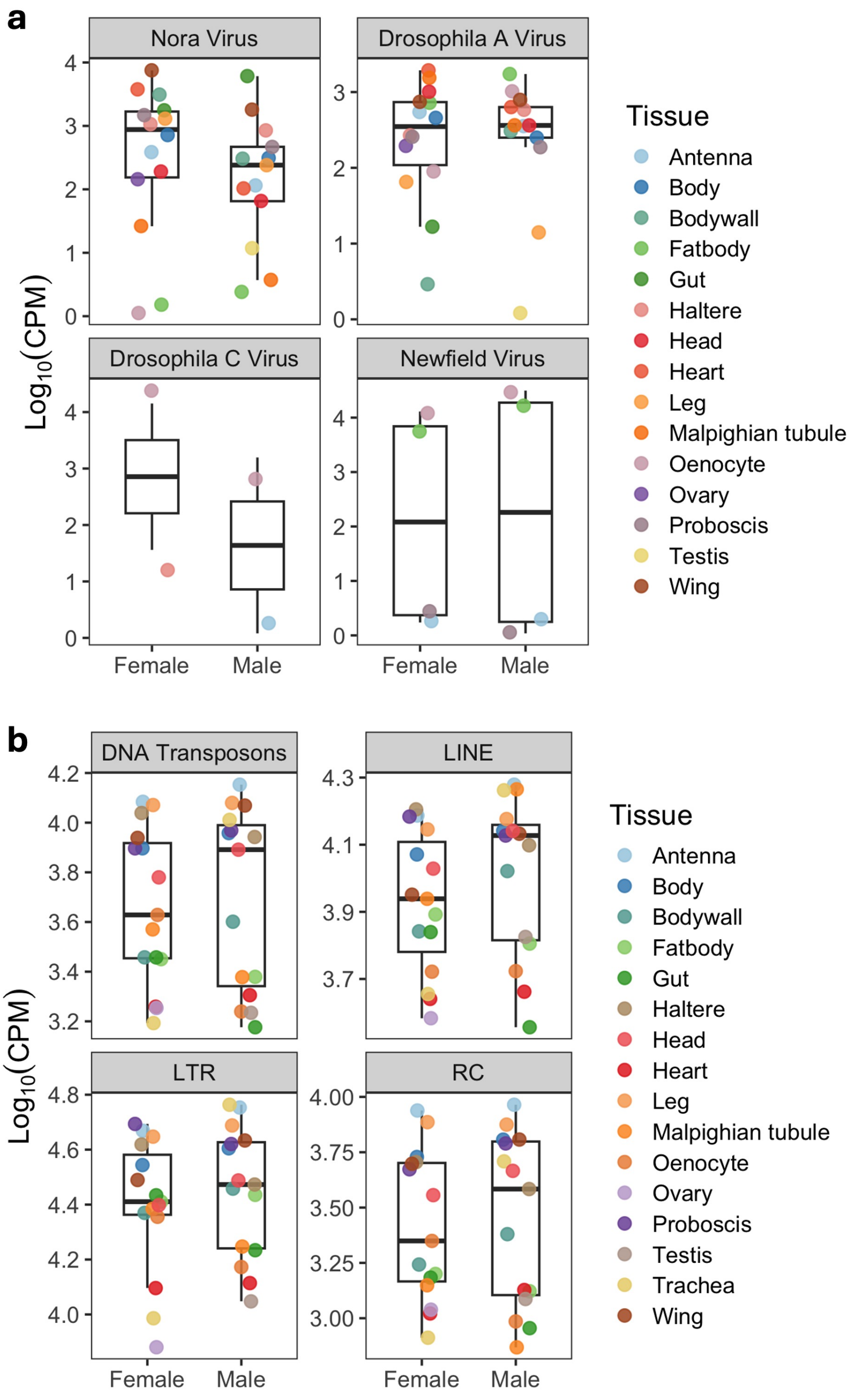


Figure S1: **Comparison of persistent RNA virus expression (viral RNA levels) and TE expression (transcripts) between male and female Drosophila.** Raw reads were normalized to counts per million (CPM). CPM values were then $\log_{10}$ transformed. Each dot indicates counts from a different tissue sample.

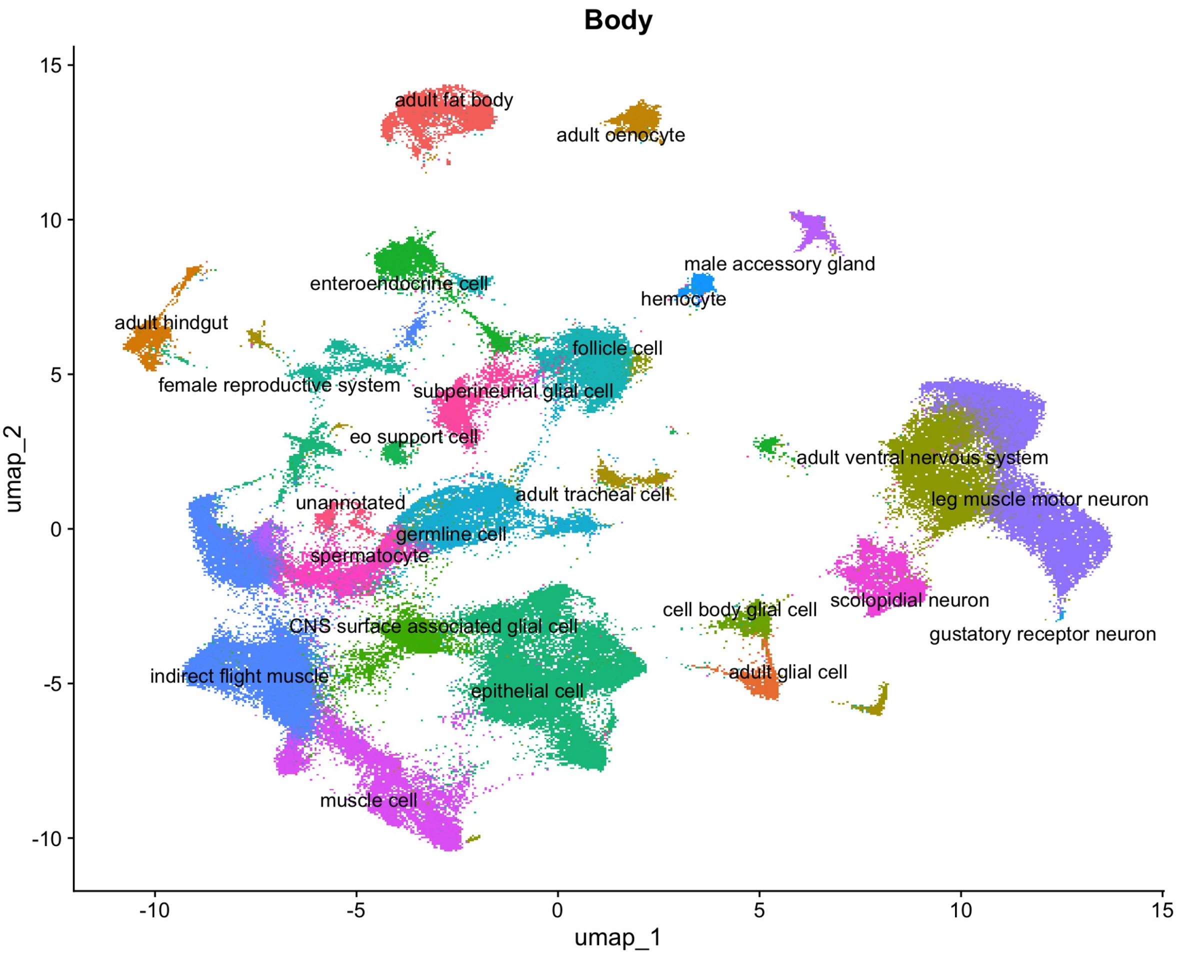


Figure S2: **Cell type landscape of the adult Drosophila whole body sample.** UMAP visualization of nuclei from the whole body dataset, colored by inferred cell types, and the annotation is based on marker gene expression (see methods). This provides the cellular context used to assess RNA virus tropism, TE expression (transcript), and infection associated transcriptional changes across major adult Drosophila cell types.

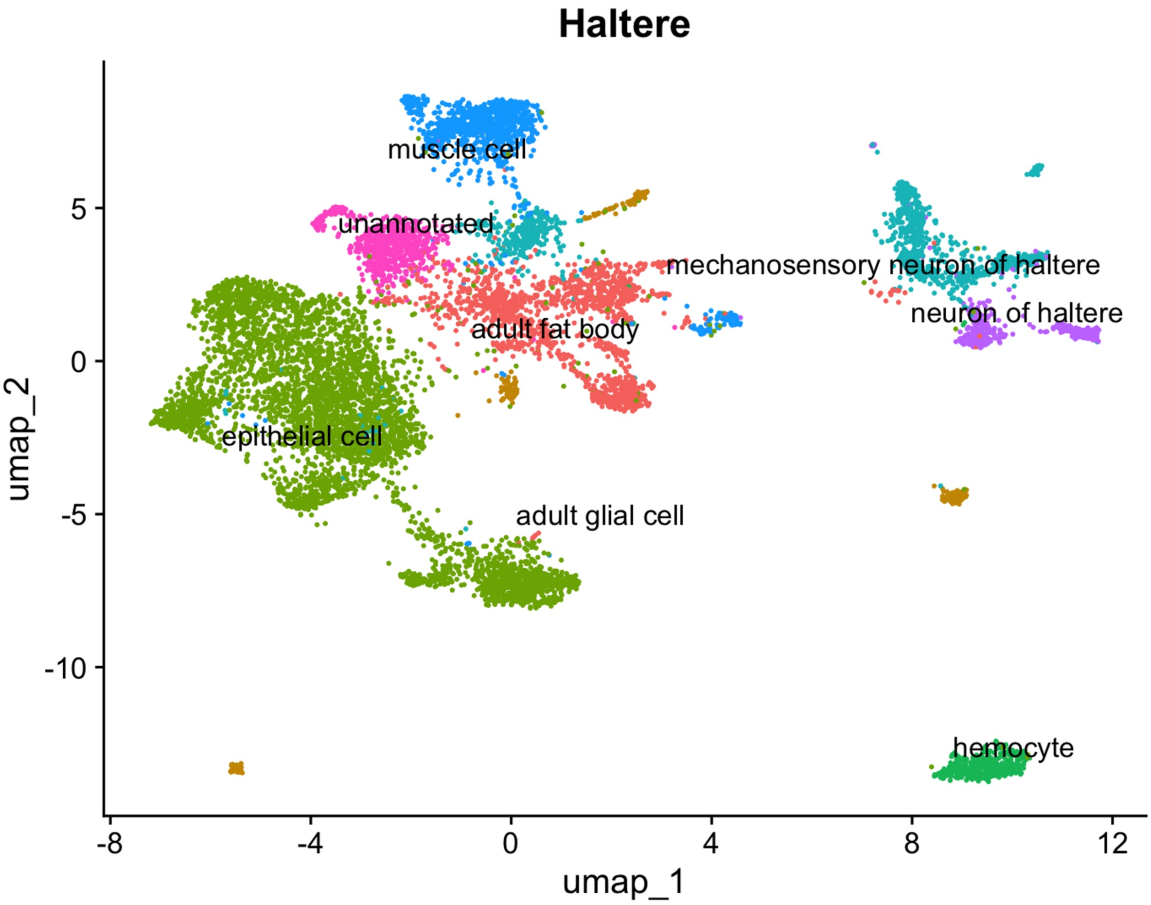


Figure S3: **Cell type landscape of the adult Drosophila haltere sample.** UMAP visualization of nuclei from the haltere dataset, colored by inferred cell types, and the annotation is based on marker gene expression (see methods).

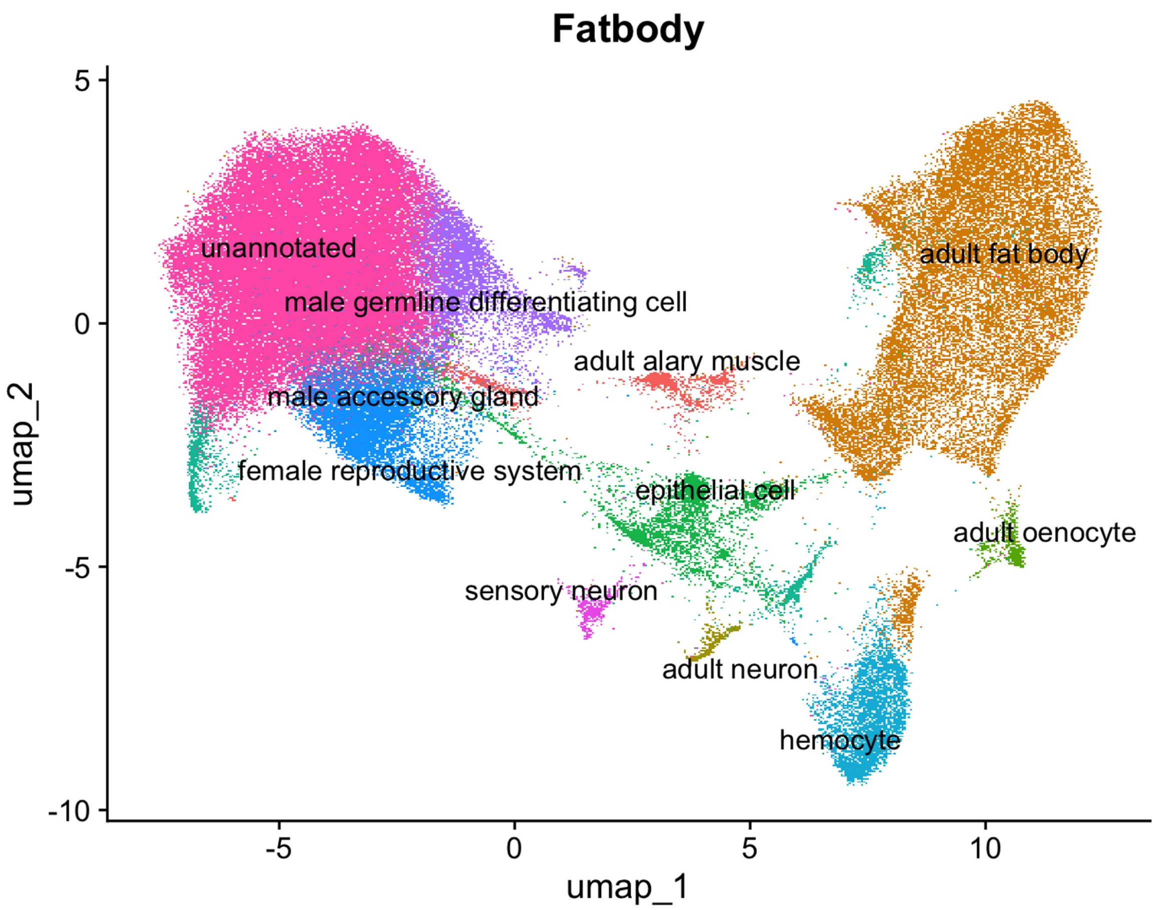


Figure S4: **Cell type landscape of the adult Drosophila fat body sample.** UMAP visualization of nuclei from the fat body dataset, colored by inferred cell types, and the annotation is based on marker gene expression (see methods).

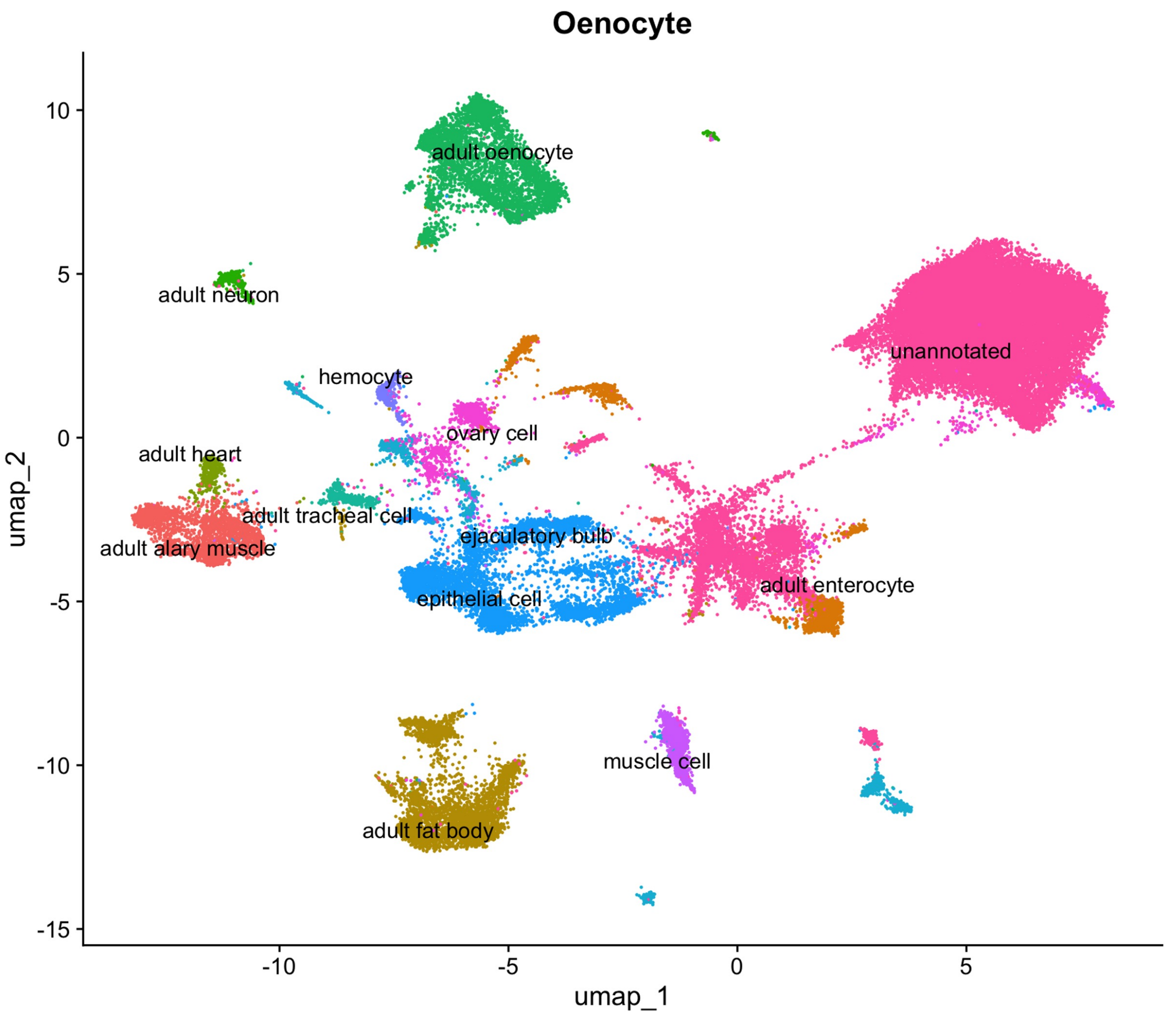


Figure S5: **Cell type landscape of the adult Drosophila oenocyte sample.** UMAP visualization of nuclei from the oenocyte dataset, colored by inferred cell types, and the annotation is based on marker gene expression (see methods).

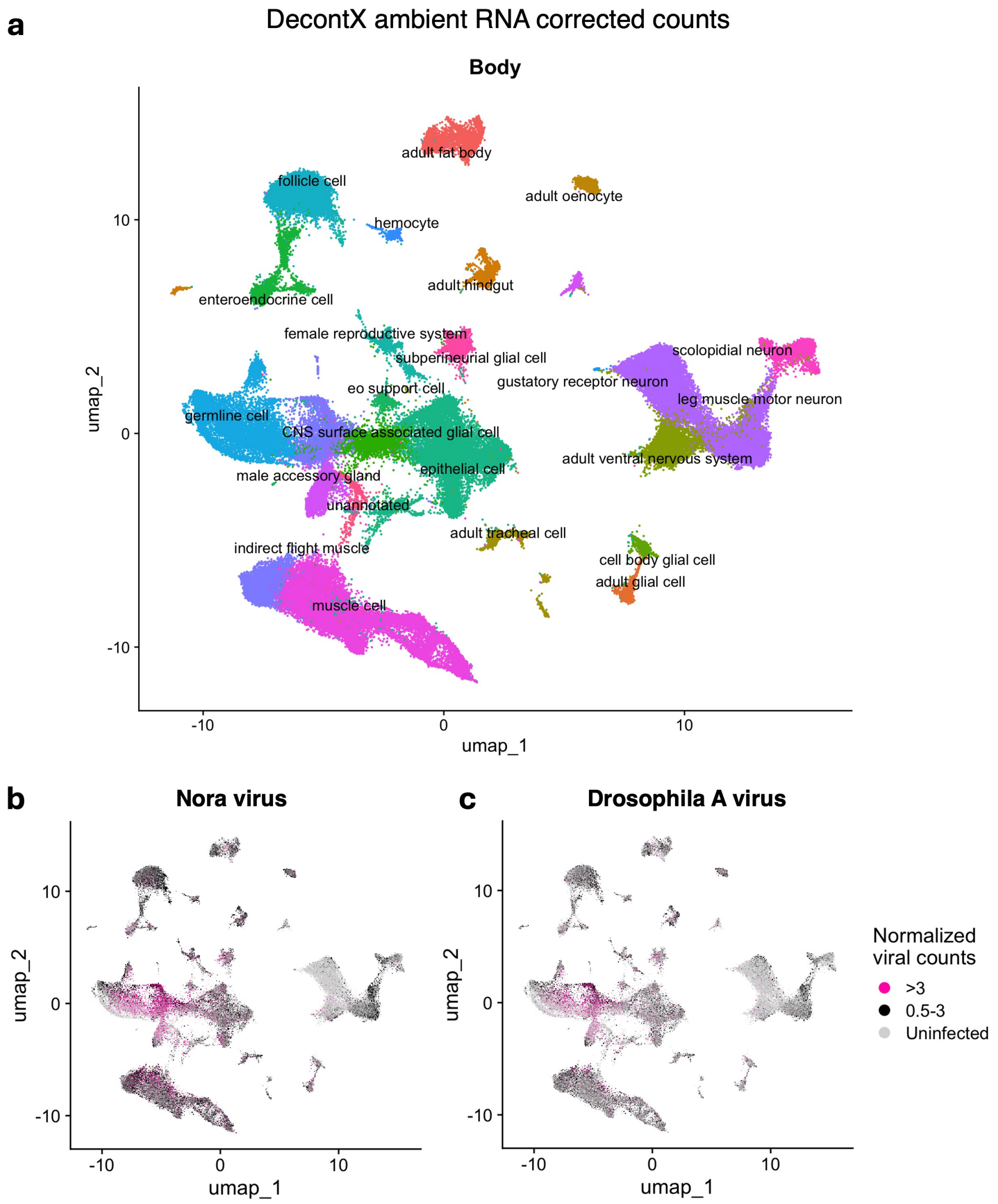


Figure S6: **Viral tropism of decontX ambient RNA corrected counts of the whole body sample of Drosophila.** (a) Cell type landscape of the adult Drosophila whole body sample. UMAP visualization of nuclei from the whole body dataset, colored by inferred cell types, and the annotation is based on marker gene expression (see methods). Nora (b), and Drosophila A virus (c) reads (viral RNA counts, counts are corrected with ambient RNA correction tool decontX) in different cell types. Counts are log normalized (which is the standard normalization method in UMAP based visualization for the single nucleus data). The Seurat LogNormalize method was used where feature counts for each cell are divided by the total counts of that cell, multiplied by a scale factor of 10,000, and then taking natural log–transformed value using log1p. >3 represents cells where viral RNA counts are greater than 3. 0.5-3 represents cells where viral read counts are between 0.5 to 3. Uninfected represents cells that don't have any viral RNA reads. Log normalized counts can be converted to CPM using the formula $CPM = (e^{\text{LogNormalized count}} - 1) \times 100$. Thus, values $> 3$ correspond to viral RNA reads greater than ∼2000 CPM, while values between 0.5 and 3 correspond to viral RNA reads between ∼65–2000 CPM.

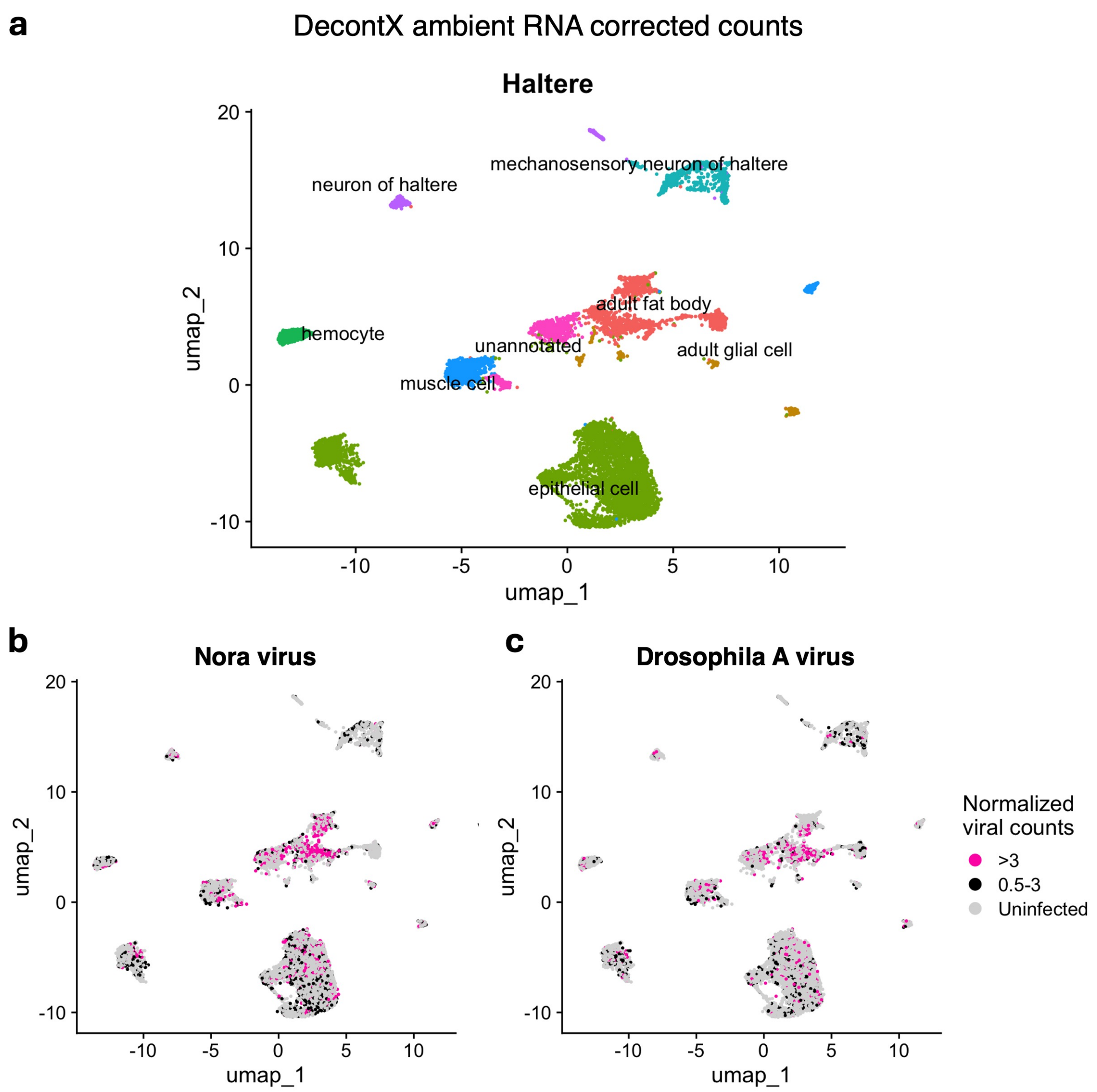


Figure S7: **Viral tropism of decontX ambient RNA corrected counts of the haltere sample of Drosophila.** (a) Cell type landscape of the adult Drosophila haltere sample. UMAP visualization of nuclei from the haltere dataset, colored by inferred cell types, and the annotation is based on marker gene expression (see methods). Nora (b), and Drosophila A virus (c) reads (viral RNA counts, counts are corrected with ambient RNA correction tool decontX) in different cell types. Counts are log normalized (which is the standard normalization method in UMAP based visualization for the single nucleus data). The Seurat LogNormalize method was used where feature counts for each cell are divided by the total counts of that cell, multiplied by a scale factor of 10,000, and then taking natural log–transformed value using log1p. >3 represents cells where viral RNA counts are greater than 3. 0.5-3 represents cells where viral read counts are between 0.5 to 3. Uninfected represents cells that don't have any viral RNA reads. Log normalized counts can be converted to CPM using the formula $CPM = (e^{\text{LogNormalized count}} - 1) \times 100$. Thus, values $> 3$ correspond to viral RNA reads greater than ∼2000 CPM, while values between 0.5 and 3 correspond to viral RNA reads between ∼65–2000 CPM.

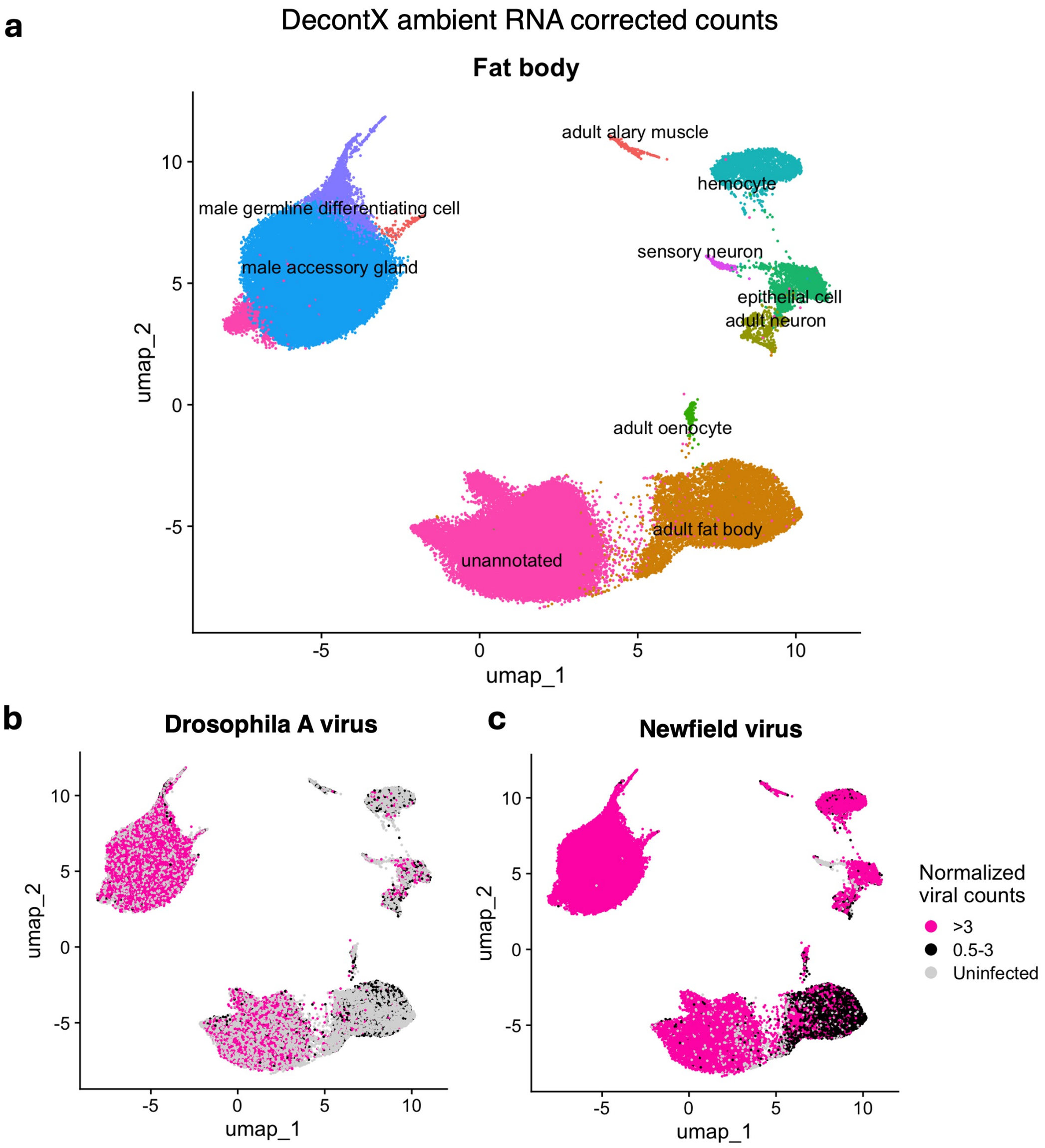


Figure S8: **Viral tropism of decontX ambient RNA corrected counts of the fat body sample of Drosophila.** (a) Cell type landscape of the adult Drosophila fat body sample. UMAP visualization of nuclei from the fat body dataset, colored by inferred cell types, and the annotation is based on marker gene expression (see methods). Drosophila A (b), and Newfield virus (c) reads (viral RNA counts, counts are corrected with ambient RNA correction tool decontX) in different cell types. Counts are log normalized (which is the standard normalization method in UMAP based visualization for the single nucleus data). The Seurat LogNormalize method was used where feature counts for each cell are divided by the total counts of that cell, multiplied by a scale factor of 10,000, and then taking natural log–transformed value using log1p. >3 represents cells where viral RNA counts are greater than 3. 0.5-3 represents cells where viral read counts are between 0.5 to 3. Uninfected represents cells that don't have any viral RNA reads. Log normalized counts can be converted to CPM using the formula $CPM = (e^{\text{LogNormalized count}} - 1) \times 100$. Thus, values $> 3$ correspond to viral RNA reads greater than ~2000 CPM, while values between 0.5 and 3 correspond to viral RNA reads between ~65–2000 CPM.

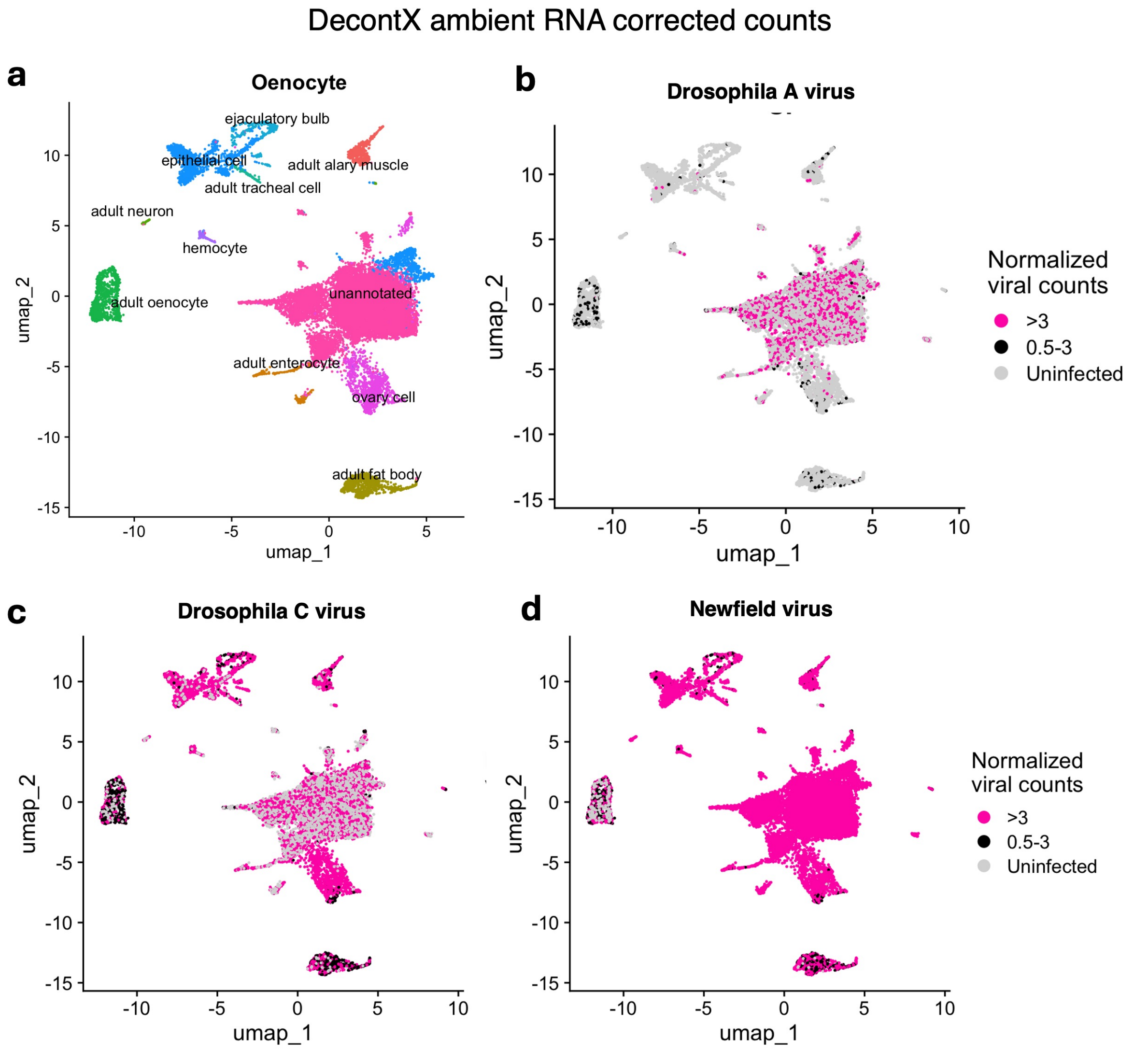


Figure S9: **Viral tropism of decontX ambient RNA corrected counts of the oenocyte sample of Drosophila.** (a) Cell type landscape of the adult Drosophila oenocyte sample. UMAP visualization of nuclei from the oenocyte dataset, colored by inferred cell types, and the annotation is based on marker gene expression (see methods). Drosophila A (b), Drosophila C (c), and Newfield virus (d) reads (viral RNA counts, counts are corrected with ambient RNA correction tool decontX) in different cell types. Counts are log normalized (which is the standard normalization method in UMAP based visualization for the single nucleus data). The Seurat LogNormalize method was used where feature counts for each cell are divided by the total counts of that cell, multiplied by a scale factor of 10,000, and then taking natural log–transformed value using log1p. >3 represents cells where viral RNA counts are greater than 3. 0.5-3 represents cells where viral read counts are between 0.5 to 3. Uninfected represents cells that don't have any viral RNA reads. Log normalized counts can be converted to CPM using the formula $CPM = (e^{\text{LogNormalized count}} - 1) \times 100$. Thus, values $> 3$ correspond to viral RNA reads greater than ∼2000 CPM, while values between 0.5 and 3 correspond to viral RNA reads between ∼65–2000 CPM.

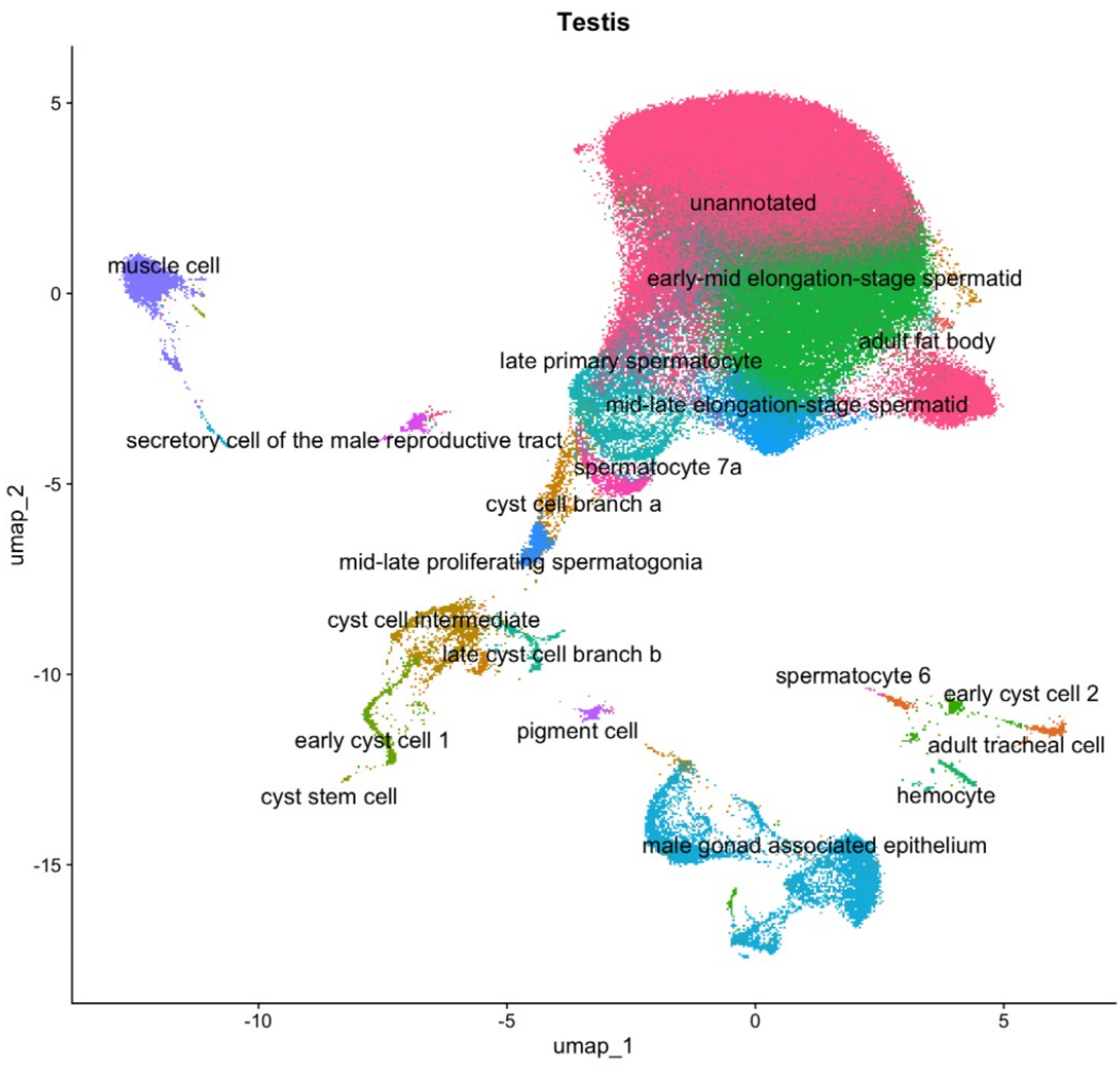


Figure S10: **Cell type landscape of the adult Drosophila testis sample.** UMAP visualization of nuclei from the testis dataset, colored by inferred cell types, and the annotation is based on marker gene expression (see methods).

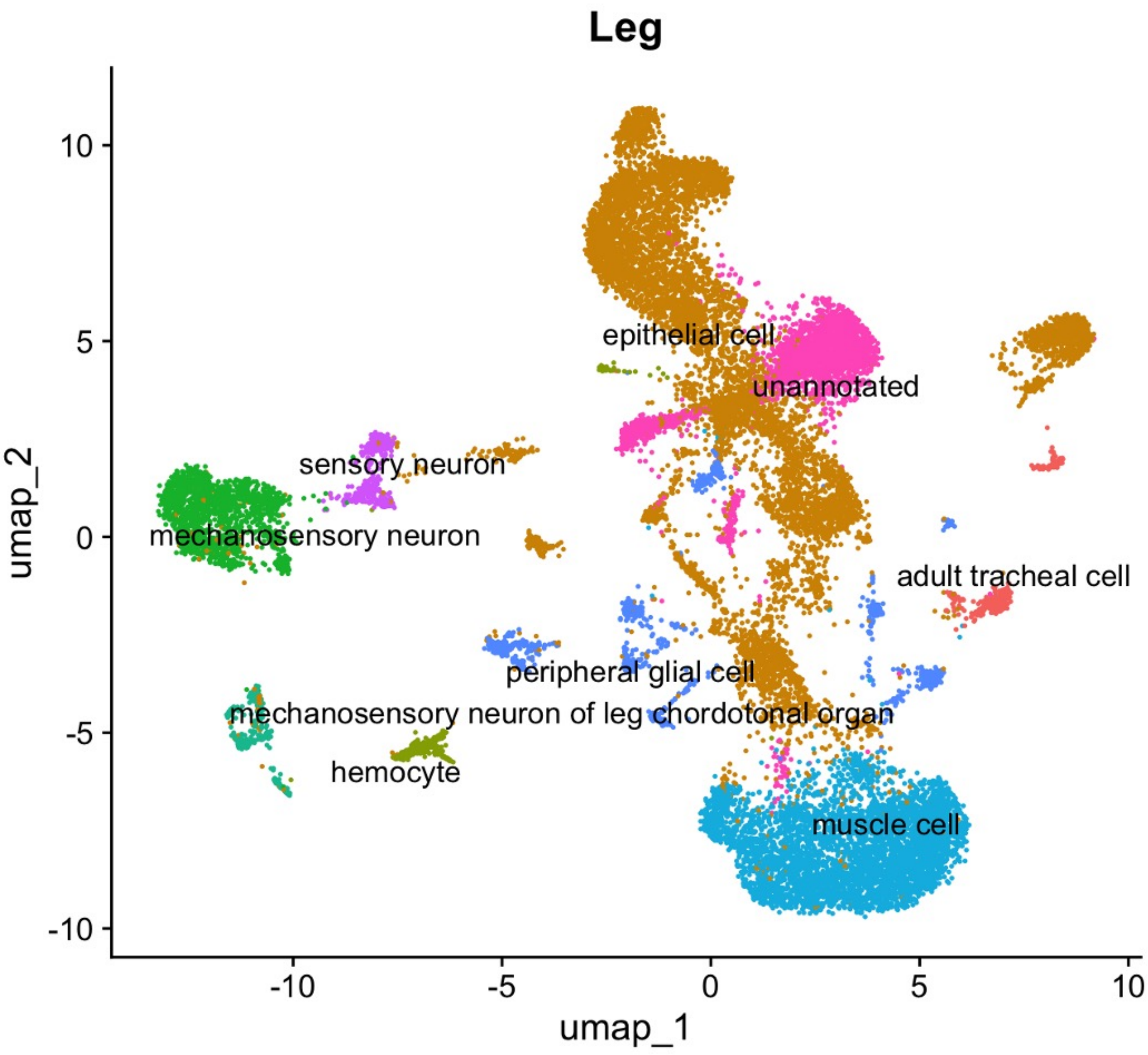


Figure S11: **Cell type landscape of the adult Drosophila leg sample.** UMAP visualization of nuclei from the leg dataset, colored by inferred cell types, and the annotation is based on marker gene expression (see methods).

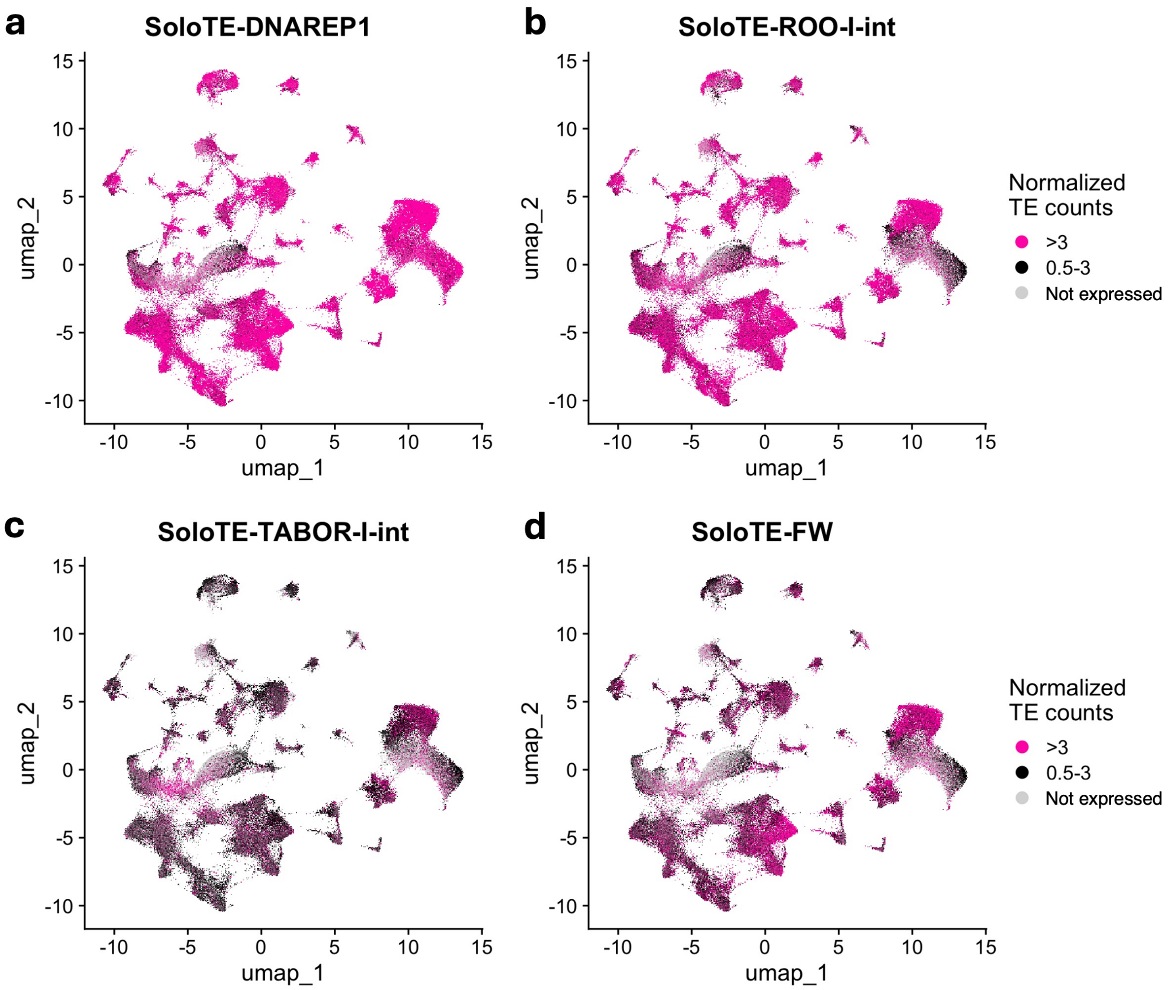


Figure S12: **Broad cellular tropism of TEs in Drosophila whole body.** UMAP plots were generated to visualize TE transcripts of DNAREP1 (Helitron:RC) (a), ROO-I-int (Pao:LTR) (b), TABOR-I-int (mdg4;Gypsy:LTR) (c), and FW (Jockey:LINE) (d) from the whole body sample. SoloTE is the tool used to measure TE transcript. Counts are log normalized (which is the standard normalization method in UMAP based visualization for the single nucleus data), and represents TE expression. The Seurat LogNormalize method was used where feature counts for each cell are divided by the total counts of that cell, multiplied by a scale factor of 10,000, and then taking natural log–transformed value using log1p. >3 represents cells where TE expression counts are greater than 3. 0.5-3 represents cells where TE expression counts are between 0.5 to 3. Not expressed represents cells that don't have any TE expression. Log normalized counts can be converted to CPM using the formula $CPM = (e^{\text{LogNormalized count}} - 1) \times 100$. Thus, values $> 3$ correspond to TE expression greater than ∼2000 CPM, while values between 0.5 and 3 correspond to TE expression between ∼65–2000 CPM.

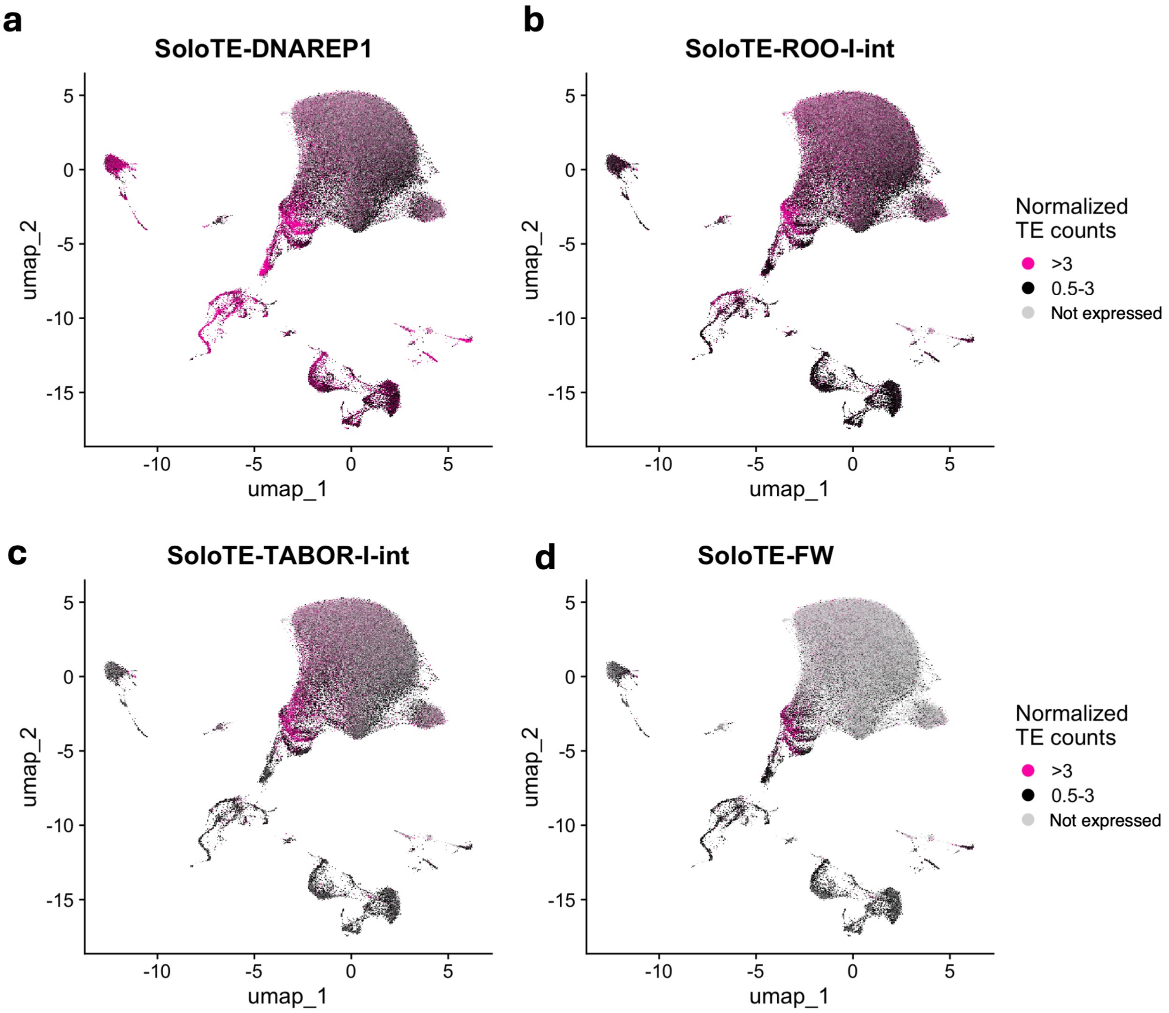


Figure S13: **Broad cellular tropism of TEs in Drosophila testis (germline tissue).** UMAP plots were generated to visualize TE transcripts of DNAREP1 (Helitron:RC) (a), ROO-I-int (Pao:LTR) (b), TABOR-I-int (mdg4;Gypsy:LTR) (c), and FW (Jockey:LINE) (d) from the testis sample. SoloTE is the tool used to measure TE transcript. Counts are log normalized (which is the standard normalization method in UMAP based visualization for the single nucleus data), and represents TE expression. The Seurat LogNormalize method was used where feature counts for each cell are divided by the total counts of that cell, multiplied by a scale factor of 10,000, and then taking natural log–transformed value using log1p. >3 represents cells where TE expression counts are greater than 3. 0.5-3 represents cells where TE expression counts are between 0.5 to 3. Not expressed represents cells that don't have any TE expression. Log normalized counts can be converted to CPM using the formula $CPM = (e^{\text{LogNormalized count}} - 1) \times 100$. Thus, values $> 3$ correspond to TE expression greater than ∼2000 CPM, while values between 0.5 and 3 correspond to TE expression between ∼65–2000 CPM.

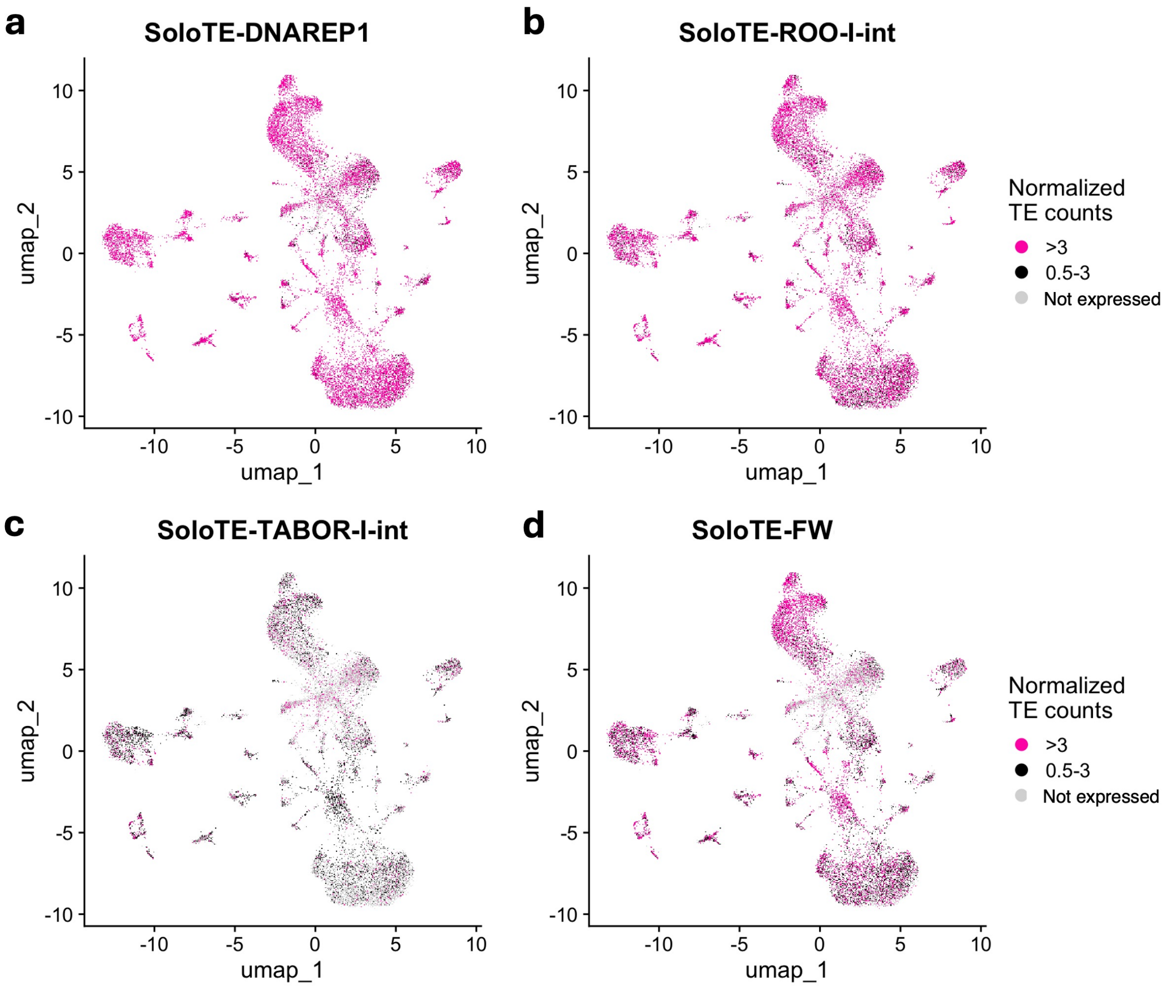


Figure S14: **Broad cellular tropism of TEs in Drosophila leg (somatic tissue).** UMAP plots were generated to visualize TE transcripts of DNAREP1 (Helitron:RC) (a), ROO-I-int (Pao:LTR) (b), TABOR-I-int (mdg4;Gypsy:LTR) (c), and FW (Jockey:LINE) (d) from the leg sample. SoloTE is the tool used to measure TE transcript. Counts are log normalized (which is the standard normalization method in UMAP based visualization for the single nucleus data), and represents TE expression. The Seurat LogNormalize method was used where feature counts for each cell are divided by the total counts of that cell, multiplied by a scale factor of 10,000, and then taking natural log–transformed value using log1p. >3 represents cells where TE expression counts are greater than 3. 0.5-3 represents cells where TE expression counts are between 0.5 to 3. Not expressed represents cells that don't have any TE expression. Log normalized counts can be converted to CPM using the formula $CPM = (e^{\text{LogNormalized count}} - 1) \times 100$. Thus, values $> 3$ correspond to TE expression greater than ∼2000 CPM, while values between 0.5 and 3 correspond to TE expression between ∼65–2000 CPM.

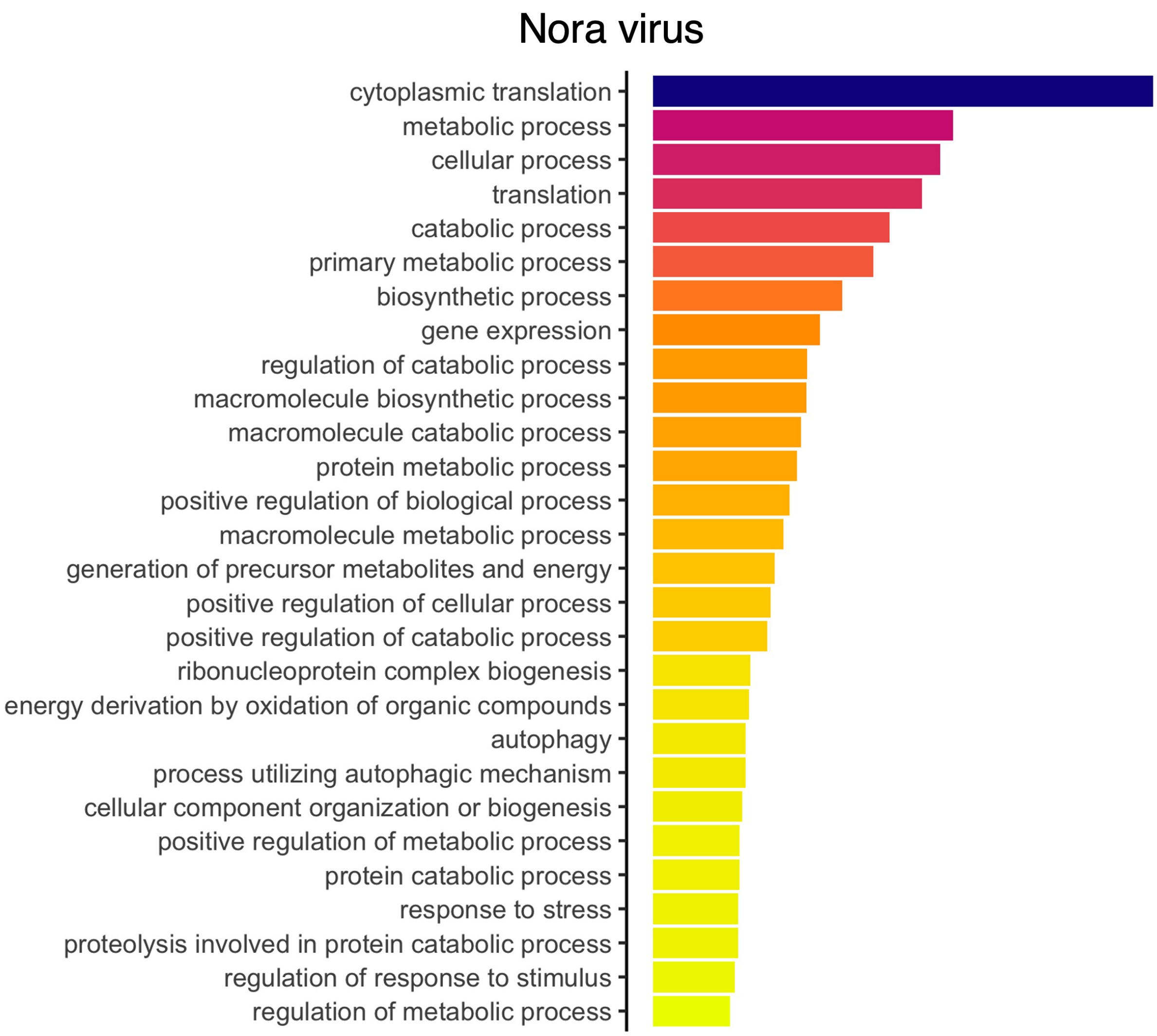


Figure S15: **Gene ontology (GO) enrichment analysis of differentially expressed genes in Nora virus infection in the whole body sample.** Differentially expressed gene set was selected using p-adjusted value $< 0.05$.

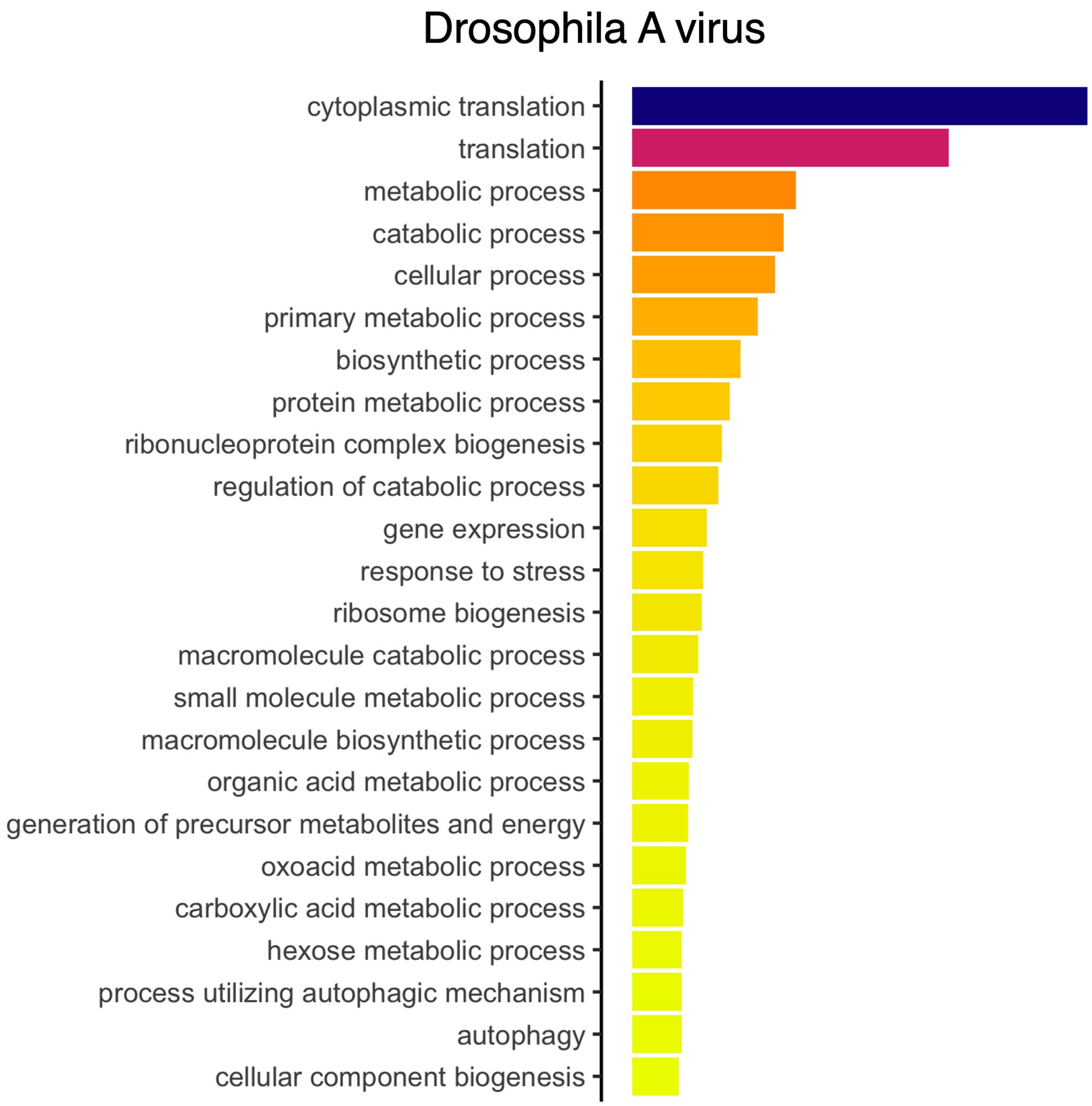


Figure S16: **Gene ontology (GO) enrichment analysis of differentially expressed genes in Drosophila A virus infection in the whole body sample.** Differentially expressed gene set was selected using p-adjusted value $< 0.05$.

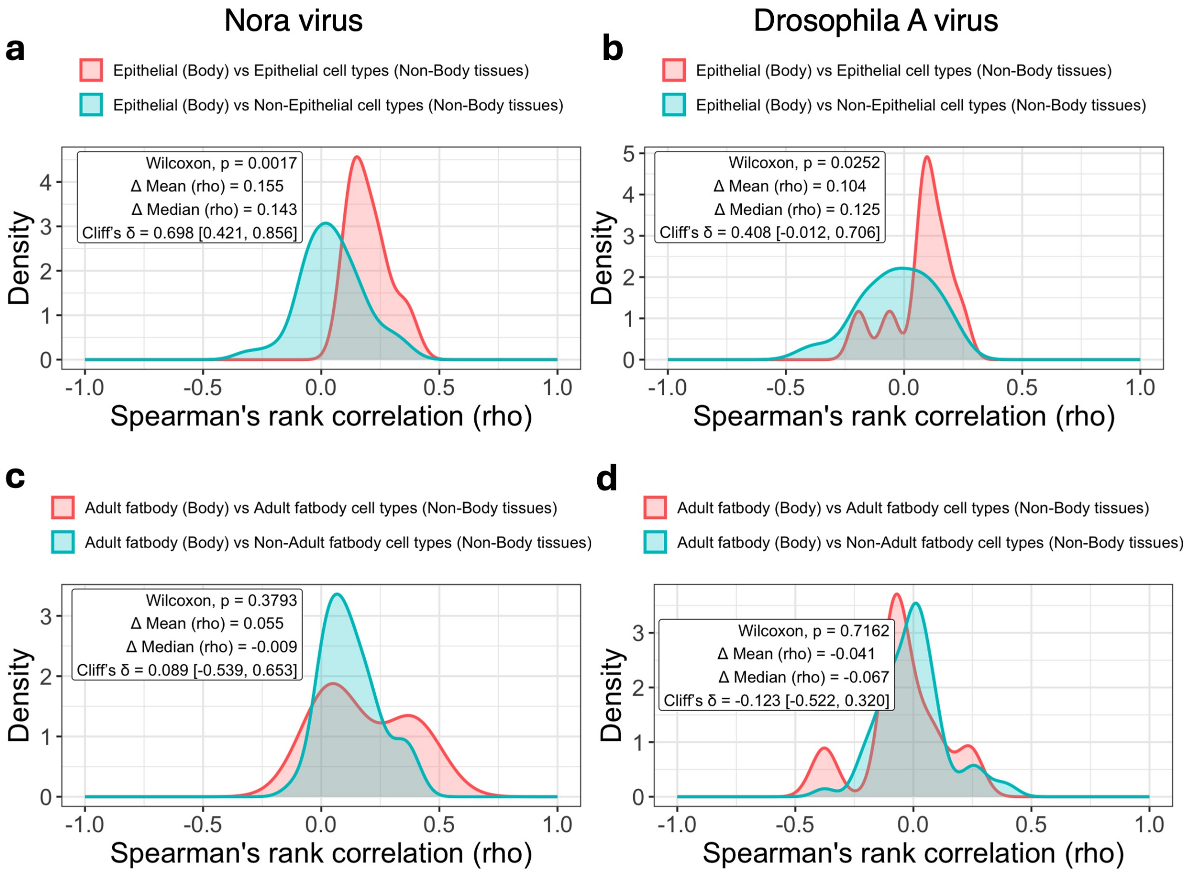


Figure S17: **Comparison of differentially expressed genes (DEGs) in similar and non-similar cell types between whole body and individual dissected tissues during viral infection.** To determine whether viral infection elicits conserved transcriptional responses within the same cell type across tissues, we compared DEGs identified in whole body cell types with those from corresponding cell types in dissected tissues. Spearman's rank correlation coefficients (rho) were calculated using $\log_2$ fold changes of DEGs from whole body epithelial cells or adult fat body cells as reference cell type and compared with the corresponding $\log_2$FC values from cell types in dissected tissues. Correlation distributions were then generated for matched cell-type comparisons (e.g., epithelial versus epithelial cell) and non-matched cell-type comparisons (e.g., epithelial versus non-epithelial cells). (a,b) Correlation distributions for Nora virus and *Drosophila A virus* infections, respectively, comparing whole-body epithelial cells with epithelial and non-epithelial cell types from dissected tissues. (c,d) Corresponding analyses for adult fat body cells during Nora virus and *Drosophila A virus* infections. Insets report the two-sided Wilcoxon rank-sum (Mann–Whitney) test p-value, differences in mean and median correlation coefficients ($\Delta$ Mean, and $\Delta$ Median), and Cliff's $\delta$ with 95% confidence intervals. Positive effect sizes indicate stronger correlations among matched cell types than among non-matched cell types.

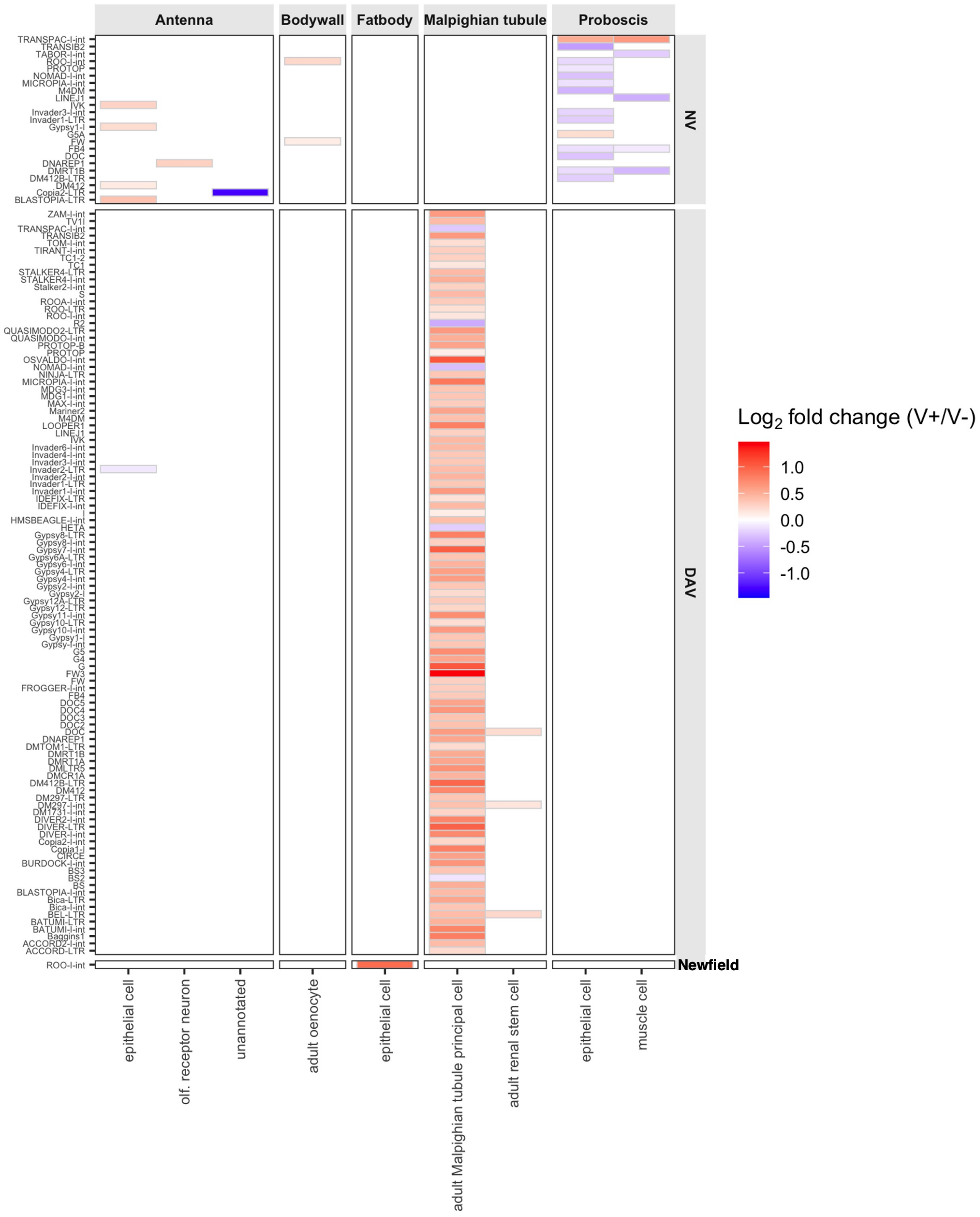


Figure S18: **TE dynamics in persistent RNA virus infectionss in different Drosophila tissues.** TE (subfamilies) expression changes during Nora virus, Drosophila A virus, and Newfield virus infection. Only tissues where significant TE subfamilies were detected are included (other tissues lacked statistical power to detect TE expression changes). Significant TE expression changes were determined using the Seurat MAST model (adjusted p-value $< 0.05$). NV = Nora virus; DAV = Drosophila A virus.

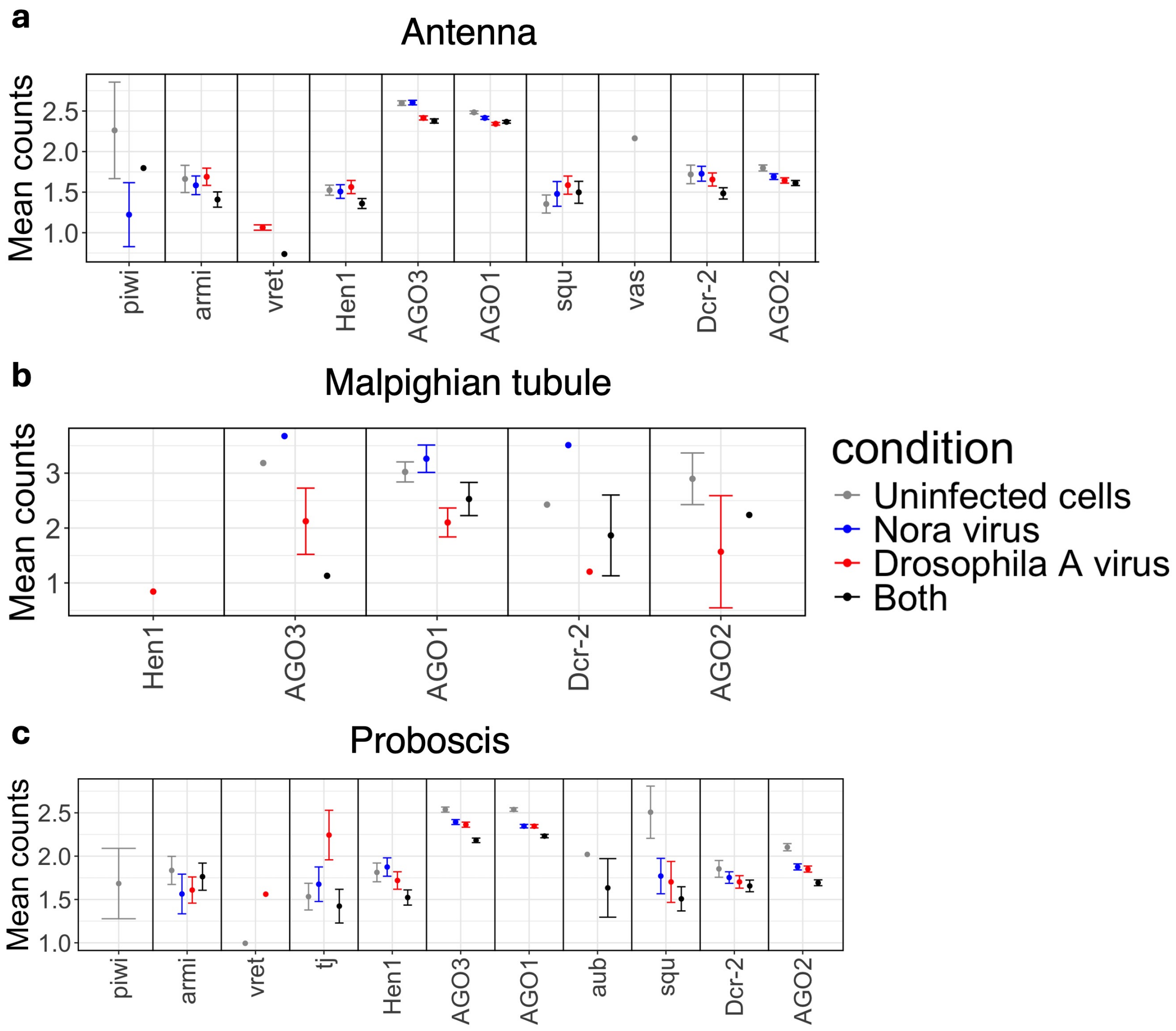


Figure S19: **Expression of piRNA and RNAi pathway genes in persistent RNA virus infections in antenna (a), malpighian tubule (b), and proboscis (c) tissue samples.** Uninfected cells are those with zero RNA counts for both Nora and Drosophila A viruses. Drosophila A virus infected cells contain Drosophila A virus RNA counts (>0) but no Nora virus counts (0). Nora virus infected cells contain Nora virus RNA counts (>0) but no Drosophila A virus counts (0). Both means cells have both Nora and Drosophila A virus RNA counts(>0). Counts are normalized with standard Seurat NormalizeData function.
